# Conditional c-MYC activation in catecholaminergic cells drives distinct neuroendocrine tumors: neuroblastoma vs somatostatinoma

**DOI:** 10.1101/2024.03.12.584622

**Authors:** Tingting Wang, Lingling Liu, Jie Fang, Hongjian Jin, Sivaraman Natarajan, Heather Sheppard, Meifen Lu, Gregory Turner, Thomas Confer, Melissa Johnson, Jeffrey Steinberg, Larry Ha, Nour Yadak, Richa Jain, David J. Picketts, Xiaotu Ma, Andrew Murphy, Andrew M. Davidoff, Evan S. Glazer, John Easton, Xiang Chen, Ruoning Wang, Jun Yang

## Abstract

The MYC proto-oncogenes (c-MYC, *MYCN*, *MYCL*) are among the most deregulated oncogenic drivers in human malignancies including high-risk neuroblastoma, 50% of which are *MYCN*-amplified. Genetically engineered mouse models (GEMMs) based on the *MYCN* transgene have greatly expanded the understanding of neuroblastoma biology and are powerful tools for testing new therapies. However, a lack of c-MYC–driven GEMMs has hampered the ability to better understand mechanisms of neuroblastoma oncogenesis and therapy development given that c-MYC is also an important driver of many high-risk neuroblastomas. In this study, we report two transgenic murine neuroendocrine models driven by conditional c-MYC induction in tyrosine hydroxylase (Th) and dopamine β-hydroxylase (Dbh)-expressing cells. c-MYC induction in Th-expressing cells leads to a preponderance of Pdx1^+^ somatostatinomas, a type of pancreatic neuroendocrine tumor (PNET), resembling human somatostatinoma with highly expressed gene signatures of δ cells and potassium channels. In contrast, c-MYC induction in Dbh-expressing cells leads to onset of neuroblastomas, showing a better transforming capacity than MYCN in a comparable C57BL/6 genetic background. The c-MYC murine neuroblastoma tumors recapitulate the pathologic and genetic features of human neuroblastoma, express GD2, and respond to anti-GD2 immunotherapy. This model also responds to DFMO, an FDA-approved inhibitor targeting ODC1, which is a known MYC transcriptional target. Thus, establishing c-MYC–overexpressing GEMMs resulted in different but related tumor types depending on the targeted cell and provide useful tools for testing immunotherapies and targeted therapies for these diseases.

## Introduction

Neuroblastoma is a solid tumor that arises from the aberrant growth of progeny of neural crest cells during early fetal development^1–3^. Tumors begin to develop via aberrant mitoses as early as the first trimester of pregnancy, with clonal evolution leading to either a favorable or an aggressive disease^2^. Neuroblastoma is the most common type of cancer in infancy and causes as much as 15% of childhood cancer mortality^4,5^. Although the outcome is excellent for children with low-or intermediate-risk disease, the 5-year survival rate for patients with high-risk neuroblastoma (principally patients older than 18 months with metastasis, or those whose tumor has MYCN amplification) remains about 50% even with intensive multimodal therapies (combined chemotherapies, stem cell transplantation, radiotherapy and anti-GD2-based immunotherapy)^5–8^. In addition, survivors of high-risk disease face long-term sequelae including hearing loss, premature aging, and subsequent malignant neoplasms due to cytotoxic chemotherapy and radiotherapy^9,10^. Unfortunately, developing safer and more effective precision therapies against high-risk neuroblastoma is challenging. One of the reasons for this is that neuroblastoma has fewer targetable recurrent mutations^11–14^. Gain-of-function mutations in the *ALK* (anaplastic lymphoma kinase) gene, with an 8% consensus mutation rate and 4% rate of amplification^15^ is the most frequently mutated oncogene currently tractable for targeted therapy in neuroblastoma^14,16–18^. Inactivating mutations in *ATRX*, which encodes a chromatin remodeling protein, were mainly identified in adolescent and young adult patients^19^ and tended to be mutually exclusive with TERT-activating alterations^20^, probably due to their redundant functions in telomere maintenance^21^. A recent pan-neuroblastoma genomic analysis further confirmed previous studies showing that the most frequently altered genes were *MYCN* (19%, primarily amplification), *TERT* (17%; structural variations), *SHANK2* (13%; structural variations), *PTPRD* (11%; structural variations and focal deletions), *ALK* (10%; somatic nucleotide variations and structural variations), and *ATRX* (8%, multiple mutation types)^11^. Genetically engineered mouse and zebrafish models of neuroblastoma have validated *MYCN* as a key oncogenic driver of neuroblastoma tumorigenesis, which can be promoted by other oncogenes such as *ALK*, *LIN28B* and *LMO1*^22–31^ (**Table 1**). These genetic models have served as valuable tools to explore and test targeted therapies and immunotherapies^32–40^, helping to advance the clinical trials of ALK inhibitors and CAR therapies^41–45^, as well as approval of DFMO by the FDA for adult and pediatric patients with high-risk neuroblastoma who have demonstrated at least a partial response to prior multiagent, multimodality therapy including anti-GD2 immunotherapy^46^. DFMO is an inhibitor of ornithine decarboxylase (ODC1) that is a direct target of MYC^47,48^, demonstrating the importance of understanding MYC biology in neuroblastoma^49^.

**Table 1.** Neuroblastoma genetic models.

| **NB GEMM and Zebrafish models** | **Genetic background** | **Note** | **References** |
| --- | --- | --- | --- |
| Th-MYCN | 129X1/SvJ | MYCN transgene | Weiss<sup>22</sup> |
| Th-MYCN | C57BL/6 | MYCN transgene | Weiss and this paper* |
| Th-MYCN/ALK<sup>F1174L</sup> | 129X1/SvJ and C57BL/6 mixed | ALK mutation introduced via transgene | Berry<sup>23</sup> |
| Th-MYCN/ALK<sup>F1178L</sup> | 129X1/SvJ and C57BL/6 mixed | ALK mutation introduced via CRISPR knockin | Van de Velde<sup>24</sup> |
| Th-MYCN/ALK<sup>Y1282S</sup> | 129X1/SvJ and C57BL/6 mixed | ALK mutation introduced via CRISPR knockin | Van de Velde<sup>24</sup> |
| Th-MYCN/ALK<sup>F1178S</sup> | 129X1/SvJ | ALK mutation introduced via homologous recombination | Borenas<sup>25</sup> |
| Th-MYCN/ALKAL2 | 129X1/SvJ | ALKAL2 introduced via transgene | Borenas<sup>25</sup> |
| Th-MYCN/Trp53<sup>KI/KI</sup> | 129X1/SvJ | Trp53 expression controlled by 4-OH-Tamoxifen | Yogev<sup>31</sup> |
| Dbh-iCre;LSL-MYCN | 129X1/SvJ and C57BL/6 mixed | Cre-mediated MYCN expression | Althoff<sup>26</sup> |
| Dbh-iCre;LSL-MYCN | C57BL/6 | Cre-mediated MYCN expression | This paper* |
| Tg(dβh:MYCN) | zebrafish | transgene | Zhu<sup>27</sup> |
| Tg(dβh:MYCN; dβh:ALK<sup>F1174L</sup>) | zebrafish | transgene | Zhu<sup>27</sup> |
| Tg(dβh:MYCN; dβh:LMO1) | zebrafish | transgene | Zhu<sup>29</sup> |
| Tg(dβh:MYCN; dβh:LIN28B) | zebrafish | transgene | Tao<sup>30</sup> |
| Tg(dβh:MYC) | zebrafish | transgene | Zimmerman<sup>57</sup> |
| Dbh-iCre;CAG-MYC | C57BL/6 | Cre-mediated MYC expression, neuroblastoma | This paper* |
| Dbh-Cre;CAG-MYC | C57BL/6 | Cre-mediated MYC expression, no tumorigenesis | This paper |
| Th-Cre;CAG-MYC | C57BL/6 | Cre-mediated MYC expression, neuroendocrine tumor, somatostatinoma | This paper |

\* indicates syngeneic models in C57BL/6 mice successfully generated in this study.

The MYC proto-oncogenes (*c-MYC*, *MYCN*, *MYCL*) encode a class of basic helix-loop-helix nuclear proteins that are altered in many cancer types^50–52^. As a transcriptional factor, MYC heterodimerizes with MAX, binds to the E box DNA consensus sequence and regulates the transcription of specific target genes that are involved in protein synthesis, metabolism, cell cycle and apoptosis^51,53–55^. MYC activation is a hallmark of cancer initiation and maintenance^56^. Normal cells can be transformed by deregulated MYC activation resulting from genomic amplification of MYC, chromosomal translocations that juxtapose MYC to enhancers for aberrant transcription, or somatic mutations leading to MYC protein stabilization. All three mechanisms of MYC activation have been identified in neuroblastoma, with 50% high-risk neuroblastomas bearing *MYCN* amplification. In contrast, a subset of high-risk neuroblastomas expresses high levels of *c-MYC* by focal enhancer amplification or genomic rearrangements leading to enhancer hijacking^57^. Pro44Leu mutation, which protects MYCN from proteasomal degradation^58^, was found in a small number of neuroblastomas^12^. Currently, neuroblastoma GEMMs are all generated based on *MYCN* oncogene expression. While these *MYCN*–driven GEMMs are invaluable for understanding the mechanism of neuroblastoma drivers and to test and develop therapies, the lack of c-MYC–driven GEMMs leaves a void in understanding neuroblastoma biology, progression, and therapeutic options as c-MYC is also a known and important driver of neuroblastomas^57^. Illuminated by a recent c-MYC–driven zebrafish neuroblastoma model^57^, this study attempted to generate c-MYC– driven GEMM. By using different Cre-lineages (tyrosine hydroxylase and dopamine β-hydroxylase) that labeled catecholamine-producing cells, we successfully developed a neuroblastoma model with high penetrance when c-MYC is activated in dopamine β-hydroxylase (Dbh-iCre) expressing cells. This model recapitulates human neuroblastoma, and is suitable for syngeneic implantation and immunotherapy testing. In contrast, somatostatinoma-like pancreatic neuroendocrine tumors occur when c-MYC is activated in tyrosine hydroxylase (Th-Cre) expressing cells, suggesting that spatiotemporal activation of the same oncogene could lead to distinct tumor entities. These new genetic models provide additional genetic tools for understanding tumorigenesis and exploring targeted therapies and immunotherapies.

## Results

### Conditional c-MYC induction by Cre recombinase under Th promoter leads to the development of pancreatic neuroendocrine tumors

We have recently generated a liver cancer model in a C57BL/6J genetic background by crossing hepatocyte-specific transgenic *Alb-Cre* mice^59^ with *CAG-STOP^flox/flox^-MYC* mice (CAG promoter-driven human c-MYC, whose expression is prevented by a LoxP site flanked STOP cassette^60^). The resultant mice showed rapid hepatic tumorigenesis with 100% penetrance after Cre-mediated activation of c-MYC^61^. We, therefore, reasoned that c-MYC activation by lineage-specific Cre recombinase may lead to tumorigenesis in any specific cell type with Cre expression. Recent scRNA-seq analysis of human adrenal medulla and neuroblastomas indicate that neuroblasts and chromaffin cells differentiated from the Schwann cell precursors (SCP) could be the cell of origin of neuroblastoma^62–65^. SCP signatures are expressed at a higher level in non-*MYCN* amplified neuroblastomas^62^, while *MYCN* and *c-MYC* expression is mutually exclusive in neuroblastomas (**Supplementary** Fig. 1a-c). SRY-Box Transcription Factor 10 (*Sox10*) is an SCP identity gene^66^, and plays an important role in neural crest and peripheral nervous system development. scRNA-seq analysis revealed that *Sox10* is highly expressed in SCP cells in comparison to its progeny, neuroblasts and chromaffin cells that express high levels of *Th* and *Dbh* instead (**Supplementary** Fig. 2a-c). We explored if activation of c-MYC in *Sox10^+^* cells could lead to neuroblastoma development by crossings of CAG-STOP^flox/flox^-MYC mice (*CAG-MYC)* with *Sox10-Cre* mice. Out of 76 mice, only 6 offspring mice were transgenic double positive, which did not match the Mendelian ratio, suggestive of embryonic lethality. Moreover, two out of 6 had brain lesions seen with magnetic resonance imaging following at around 270 days (**Supplementary** Fig. 3). These data suggest that *Sox10-Cre* may not be suitable for generating neuroblastoma models.

Previous studies have shown that *MYCN* activation in either tyrosine hydroxylase (Th) or dopamine β-hydroxylase (Dbh)-positive cells led to the development of neuroblastomas^22,26^, while c-MYC induction by *Dbh* promoter in zebrafish also resulted in neuroblastoma occurrence^57^. We therefore surmised that activation of c-MYC by breeding *CAG-MYC* mice with *Th-Cre* or *Dbh-Cre* may also lead to the transformation of sympathetic neurons into neuroblastoma. To test this, we first bred a *Dbh-Cre* knock-in (*Dbh^Cre^KI*) mouse strain with the *CAG-MYC* mouse. *Dbh^Cre^KI* expresses Cre recombinase from the endogenous *Dbh* locus, abolishing endogenous *Dbh* gene expression^67^. However, no tumorigenesis was observed in the compound mice routinely monitored by ultrasound imaging examination. Then, we utilized a *Th-Cre* mouse strain to breed the *CAG-MYC* mice. The *Th-Cre* transgenic mice have a rat *Th* gene promoter directing Cre recombinase expression in catecholaminergic cells^68^. Over a 600-day follow-up of the *Th-Cre;CAG-MYC* mice, we observed that 30% of mice developed abdominal tumors (12 out of 43 mice, **Fig. 1a-c**), while only 0.6% of the control mice developed tumors (1 out of 16 mice, **Fig. 1b**), consistent with spontaneously arising, age-related neoplasia. The tumor growth rate of *Th-Cre* mice also seemed to be slower (**Fig. 1c**), so that the survival curves for *Th-Cre* mice were not statistically different from wild-type mice that died of spontaneously arising and age-related background lesions (**Fig. 1c, 1d**). Surprisingly, necropsy and histopathology examination of the tumor locations revealed that the tumors observed by ultrasound imaging in the abdomen were anatomically present in the pancreas and duodenum. The difference in anatomic location suggested that the *Th-Cre;CAG-MYC* tumors may not always be neuroblastoma since neuroblastoma mainly arises from the superior cervical ganglion, adrenals or celiac ganglion in mouse models^22,26^. There were no macroscopic tumor metastases observed in any other organs although lymphovascular invasion was observed around some grossly visible tumors. Hematoxylin and eosin (H&E) staining showed that neoplastic cells were epithelioid in morphology and small to medium in size, with fine fibrovascular stroma, which are morphologic features that are typical for neuroectodermal tumors. Often pancreatic ductal glands were entrapped in the tumors, which suggested these neoplasms may be arising from or near the pancreas and duodenum and invading into the pancreas (**Fig. 1e, 1j, HE**). Immunohistochemistry (IHC) staining of tumors showed that all neoplasms were immuno-positive for neuronal specific enolase (NSE, **Fig. 1f**) and synaptophysin (SYP, **Fig 1k**).

**Figure 1.**
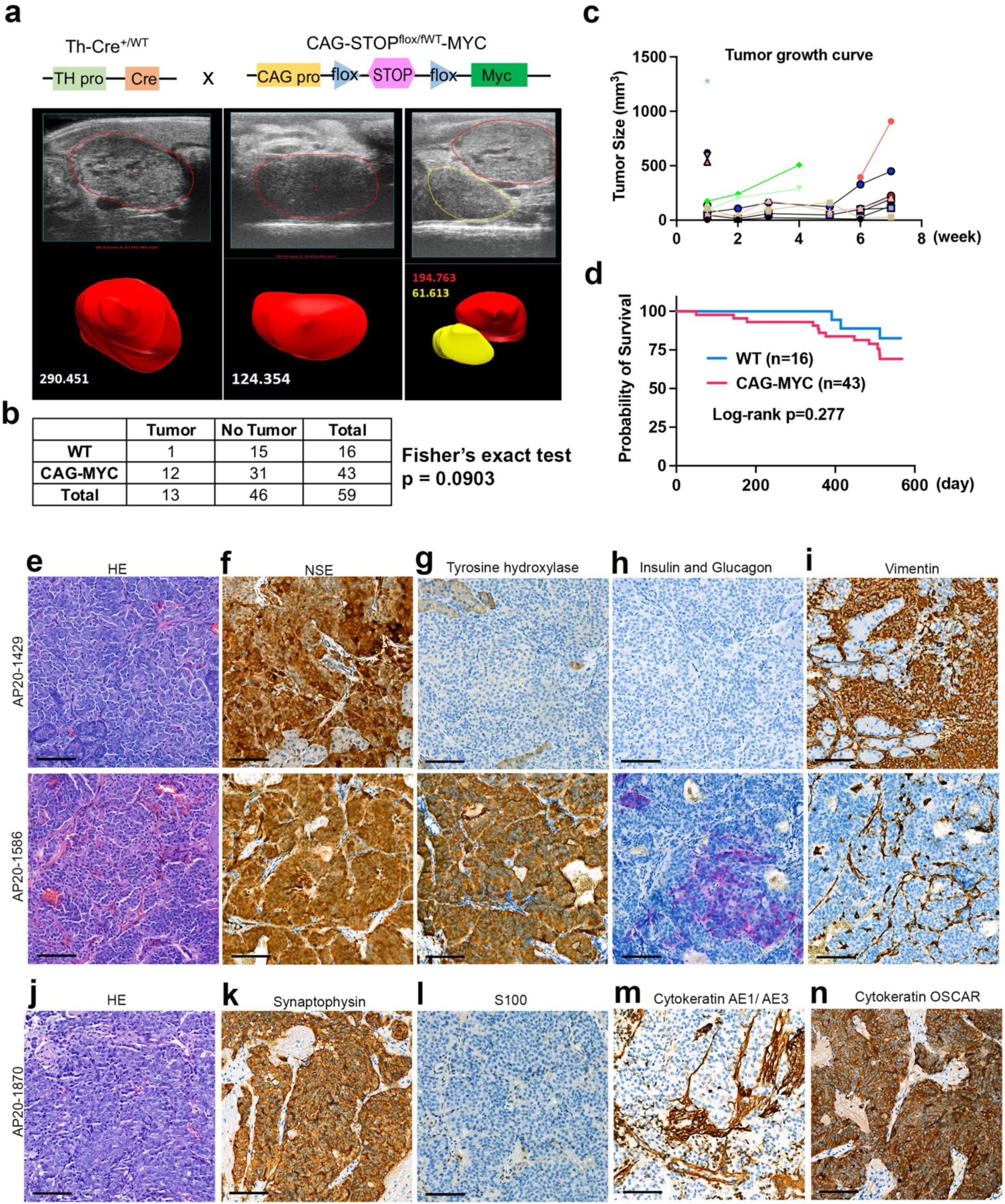
Generation of Th-Cre;CAG-MYC tumors. **a.** Ultrasound imaging of tumors in Th-Cre;CAG-MYC mice as indicated by breeding strategy. The top panel of ultrasound images are two-dimensional ultrasound images while the bottom ones are volume reconstructions. The red and yellow indicate two tumors. **b.** Numbers of Th-Cre;CAG-MYC mice with tumors. The p value is calculated by Fisher’s exact test to compare with the wildtype mice. **c.** Tumor growth curve monitored by ultrasound imaging for each mouse. **d.** Kaplan-Meier analysis of mouse survival for Th-Cre;CAG-MYC and wildtype mice. The log-rank P value is calculated by Mantel-Cox test. **e-n**. Representative HE and immunohistochemical staining of neuroendocrine tumors arising in or near the pancreas. Scale bar = 100μm.

Neuroendocrine neoplasms include well-differentiated neuroendocrine tumor (NET), poorly differentiated neuroendocrine carcinoma (NEC), pheochromocytoma (adrenal medulla), and paraganglioma (extra-adrenal). While NET and NEC tumors are epithelial origin, the pheochromocytoma and paraganglioma are non-epithelial. However, these tumors can have comparable histologies (i.e., nested architecture), expression of general neuroendocrine markers such as NSE and SYP as well as the production of peptide hormones and/or biogenic amines^69^. As both NET/NEC and paraganglioma can also occur in the pancreas, we attempted to differentiate the origin of Th-Cre;CAG-MYC tumors by examining the epithelial markers including the pan-cytokeratin markers AE1/AE3 and OSCAR. While AE1/AE3 was negative by IHC, the tumors showed immunoreactivity for OSCAR indicating that these tumors are of epithelial origin (**Fig. 1m, n**). Nevertheless, epithelial markers in clinical samples could be negative in some NET but positive in some paraganglioma. Two out of 4 tumors were positive for TH (**Fig. 1g**). One tumor showed weak glucagon staining (**Fig. 1h**) in some areas, but all tumor cells were determined to be immuno-negative for insulin suggesting that a combination of functional and non-functional NETs may arise in this model. TH could be either positive or negative in both NET/NEC and paraganglioma. Additionally, S100, another neuroendocrine marker for paraganglioma and NET was immuno-negative (**Fig. 1l**) although immunoreactivity was observed in the stroma. One tumor (**Fig 1i**, AP20-1429) was immuno-positive for vimentin. Paragangliomas may express vimentin. Additionally, a subset of human pancreatic neuroendocrine tumors and other neuroendocrine carcinomas arising in other organs may also express vimentin. The presence of vimentin expression in neuroendocrine carcinomas has been correlated with dedifferentiation, epithelial to mesenchymal transition, and increased biologically aggressive behavior such as increased invasion and/or metastatic capability^25^. These data suggest that *Th-Cre*;CAG-MYC tumors may be neuroendocrine tumors originating from islet cells, although the possibility of a paraganglioma originating from the sympathetic nervous system cannot be entirely ruled out. The pathology evaluations suggest both epithelial and non-epithelial neuroendocrine tumors may arise in this model system.

### The *Th-Cre;CAG-MYC* tumors show a remarkable induction of the potassium channel gene signature

To further characterize the molecular features of the *Th-Cre;CAG-MYC* tumors, we performed bulk RNA-seq analysis of tumors and the age-matching normal pancreatic tissues, followed by gene set enrichment analysis for hallmark gene signatures. In comparison with the normal controls that expressed high levels of gene sets including ‘oxidative phosphorylation’, ‘fatty acid metabolism’, ‘KRAS signaling down’ and ‘p53 pathway’, the tumors expressed high levels of genes signatures of ‘hedgehog signaling’, ‘E2F targets’, ‘KRAS up signaling’ and ‘epithelial-to-mesenchymal transition’ (**Supplementary** Fig. 4a**, 4b**). Then we examined the classical signaling pathways altered in tumors (**Table S1**) and found that the top gene signature upregulated in tumors was ‘potassium channels’, while the downregulated ones were ‘eukaryotic translation initiation’ and ‘respiratory chain transport’ (**Fig. 2a-f**), suggesting that tumor cells undergo metabolic reprogramming. The potassium channel genes upregulated in tumors encode ATP-sensitive potassium channels such as ABCC8 and KCNJ11 that regulate insulin secretion in pancreatic β cells. The genes encoding voltage-gated potassium channels such as KCNQ2 also play an important role in regulating pancreatic endocrine function, further suggesting that the *Th-Cre;CAG-MYC* tumors might be PNET. However, the biological impact of such a broad induction of potassium channel genes in this genetic model awaits further studies.

**Figure 2.**
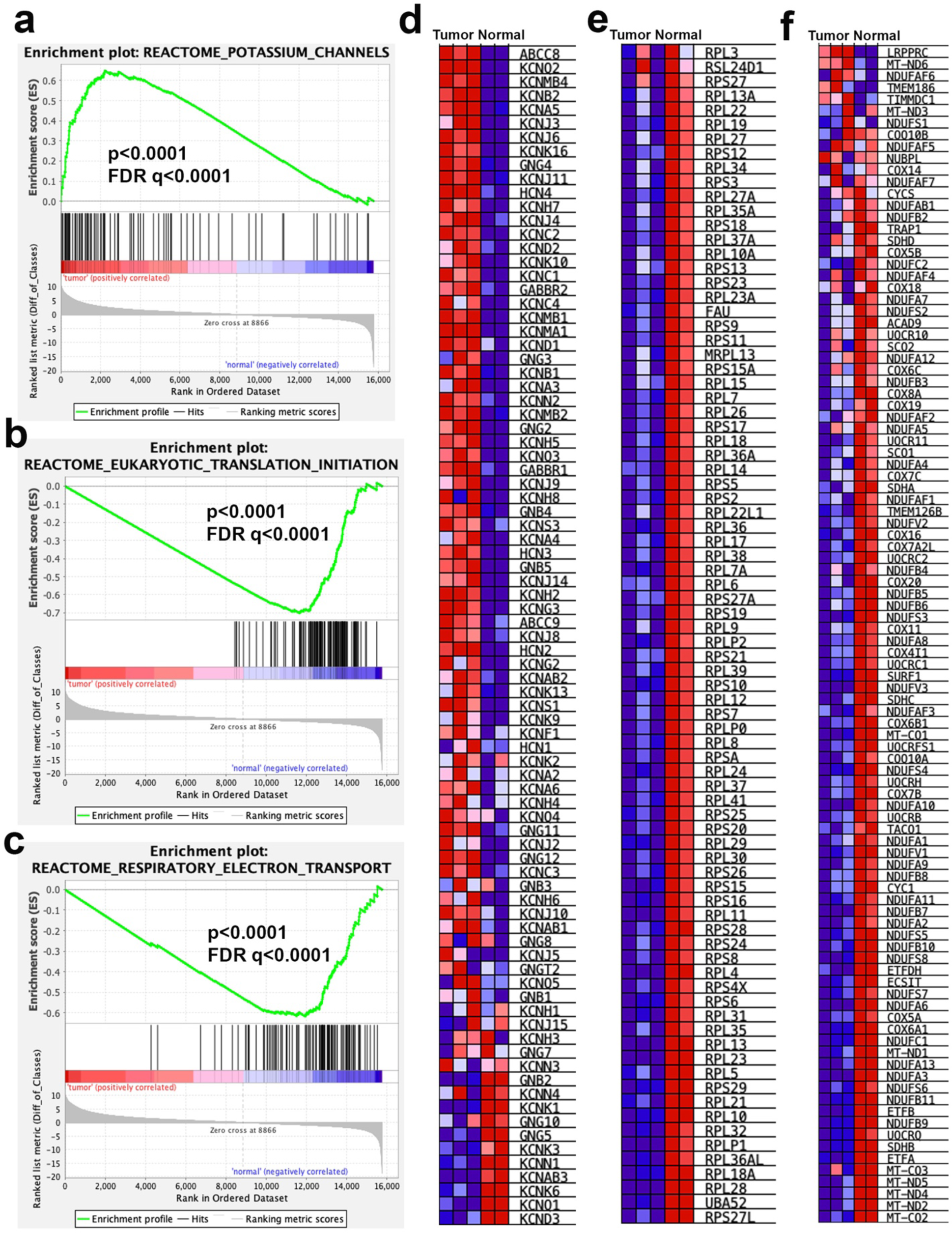
Pathway enrichment analysis of Th-Cre;CAG-MYC tumors by GSEA. **a.** GSEA result shows that genes encoding potassium channels are highly expressed in Th-Cre;CAG-MYC tumors in comparison with the pancreatic normal tissues. **b.** GSEA result shows that eukaryotic translation initiation genes are highly expressed in Th-Cre;CAG-MYC tumors in comparison with the pancreatic normal tissues. **c.** GSEA result shows that respiratory electron transport genes are highly expressed in Th-Cre;CAG-MYC tumors in comparison with the pancreatic normal tissues. **d-f**. Heatmaps for Figure 2a-c respectively.

### The *Th-Cre;CAG-MYC* tumors have molecular features resembling somatostatinoma originating from δ cell of pancreatic islets

Neuroendocrine tumors (NETs) are heterogeneous neoplasms originating from neuroendocrine system cells with traits of both hormone-producing endocrine cells and nerve cells^70^. NETs could arise in anywhere in the body, but most occur in lungs and the gastroenteropancreatic (GEP) digestive tract. The GEP-NETs represent the second most common digestive cancer^71^, which includes carcinoid tumors of the gastrointestinal tract and pancreatic NETs (PNETs). Carcinoid tumors originate from enterochromaffin cells of the gut. However, the cell of origin of PNETs is not definitive. Whereas PNETs are commonly thought to arise in the islets of Langerhans, one study proposed that the precursors in the ductal/acinar units are an alternative source of PNETs^72^. The islets of Langerhans in the pancreas are composed of hormone-producing cell types including α cells (glucagon), β cells (insulin), δ cells (somatostatin) and PP cells (pancreatic polypeptide), G cells (gastrin) and ε cells (ghrelin); while PNET can be classified as gastrinoma, insulinoma, glucagonoma and other rare types such as VIPoma and somatostatinoma based on their cell of origins or hormones they produce. To further understand the molecular features of the *Th-Cre;CAG-MYC* tumors, we performed gene set enrichment analysis of cell type signatures to determine the cell identity of tumor cells. We found that the transcriptional profiles of the normal tissues mostly resembled the gene set of fetal pancreatic acinar cells, while the tumor cells most significantly resembled the cell types of fetal pancreatic islet endocrine cells, followed by islet delta cells (**Fig. 3a, 3b**), suggesting that the *Th-Cre;CAG-MYC* tumors when arising in the pancreas are highly likely to be somatostatinoma, a rare type of PNET whose cell of origin is δ type cell. To further confirm the molecular features of this genetic model, we performed single nuclear RNA sequencing (snRNA-seq) analysis to examine the expression of representative markers of each type of PNET. The islet cells in pancreas are derived from progenitor cells directed by pancreatic and duodenal homeobox 1 (Pdx1) and other transcriptional factors such as Pax6^73^ (**Fig. 3c**). UMAP analysis of snRNA-seq data identified 6 populations of cells in one tumor mass, demonstrating the cellular heterogeneity of tumor cells (**Fig. 3d**), which expressed high levels of *Pdx1* and *Pax6* in nearly all populations (**Fig. 3e, 3f**), further confirming that pancreatic endocrine cells are the origin of tumors as the expression of PDX1 in pheochromocytoma and paraganglioma was extremely rare (**Supplementary** Fig. 5). In line with the bulk RNA-seq results, snRNA-seq showed very low or undetectable levels of *Gcg* (encoding glucagon, a marker for α cells and glucagonoma), *Ins1* (encoding insulin, a marker for β cells and insulinoma), *Gast* (encoding gastrin, a marker for G cells and gastrinoma), *Vip* (a marker for VIPoma), and Ghrl (encoding ghrelin, a marker for ε cells)^74^ (**Fig. 3g-k**), and low levels of PP cell markers *Ahr* and *Ppy* that encodes aryl hydrocarbon receptor and pancreatic polypeptide^75^, respectively (**Fig. 3l**). However, tumor cells expressed high levels of *Sst* (encoding somatostatin, a marker for δ cells and somatostatinoma)^74^, *Sstr3* (encoding somatostatin receptor 3) and *Isl1* (encoding ISL LIM homeobox 1, a δ cell transcriptional factor)^75^ (**Fig. 3m**). The general NET markers such as *Chga* (chromogranin A), *Syp* (Synaptophysin), *Cd56* (NCAM1) were highly expressed in these tumor cells, and *Pgr* (progesterone receptor), another PNET marker was also positive (**Supplementary** Fig. 6). *Gata3*, which is nearly universally positive in pheochromocytoma and paraganglioma (**Supplementary** Fig. 5), was negative in this tumor, while the epithelia marker *Epcam* was positive (**Supplementary** Fig. 6). However, *Sox10*, the marker of Schwann cell lineage malignancies, was negative (**Supplementary** Fig. 6). We had similar results from analysis of a second *Th-Cre;CAG-MYC* tumor which support the δ cell as the tumor origin (**Supplementary** Fig. 7). Taken together, these data suggest that we have generated a somatostatinoma PNET by crossing *Th-Cre* and *CAG-MYC* mice.

**Figure 3.**
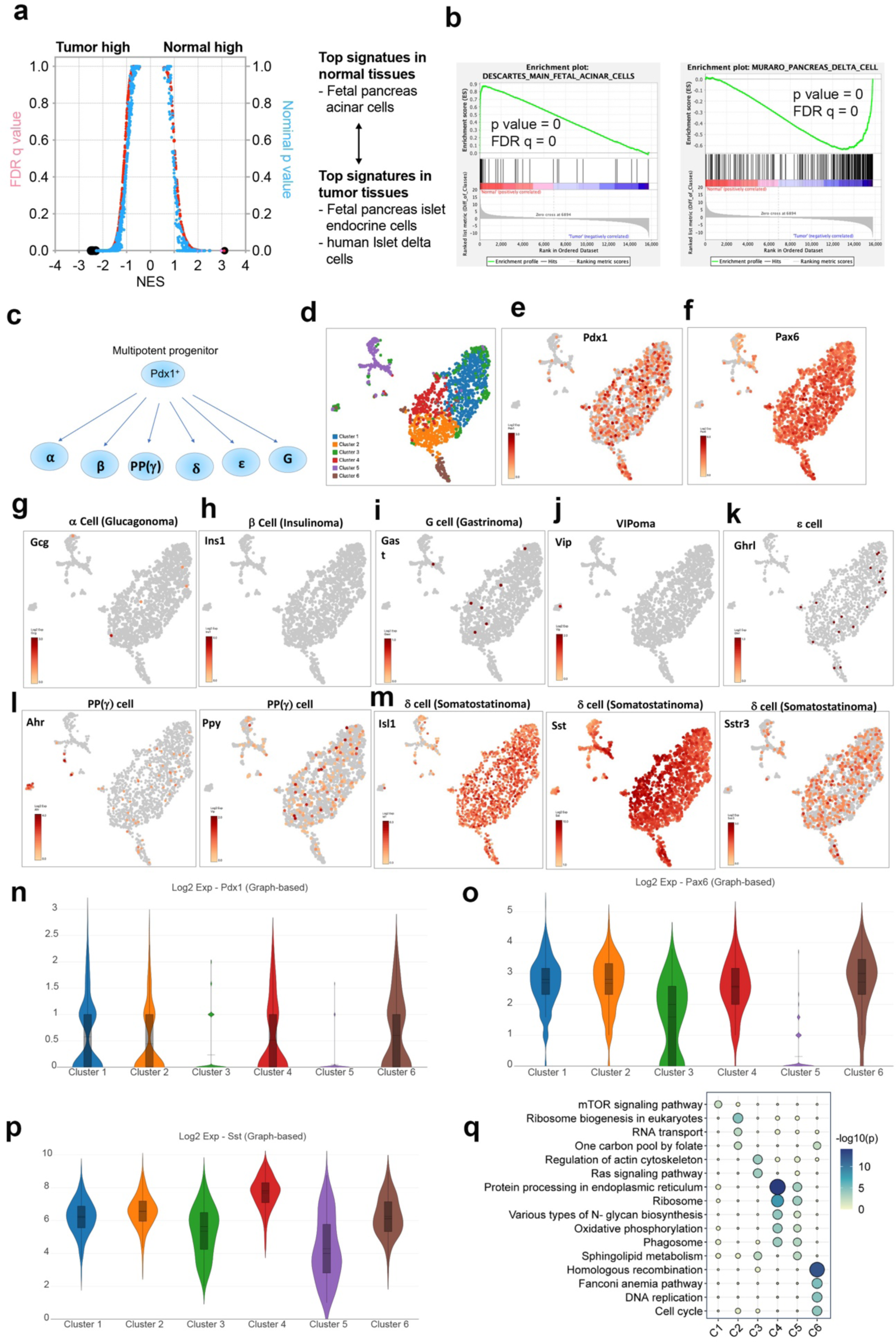
Th-Cre;CAG-MYC tumors resemble somatostatinoma originated from δ cells of pancreatic islets. **a.** Quantitative comparison of all gene sets (n = 737) from the cell type signatures by gene set enrichment analysis (GSEA) for increased (left) and reduced (right) expression of genes in Th-Cre;CAG-MYC tumors vs normal pancreatic tissues. Data are presented as a scatterplot of normalized p value/false discovery q value vs normalized enrichment score (NES) for each evaluated gene set. The gene sets circled in black color indicate ‘fetal pancreas islet endocrine cells’, ‘human islet delta cells’ (left) and ‘fetal pancreas acinar cell’ (right). **b.** GSEA results show signatures of ‘fetal acinar cell’ (left) and ‘pancreas delta cells’ (right) that are downregulated and upregulated in Th-Cre;CAG-MYC tumors. **c.** A schematic view of cell types in pancreatic islets originated from Pdx1^+^ precursors. **d.** UMAP shows 6 clusters of cells in a Th-Cre;CAG-MYC tumor by analysis of single nuclear RNA-seq. **e-m**. Respective markers for each cell type of islets or pancreatic neuroendocrine tumors. **n-p**. Violin plots show expression of Pdx1, Pax6 and Sst in each cluster of the Th-Cre;CAG-MYC tumor. **q.** Bubble plot shows pathway enrichment of the top 100 genes in each cluster. The top differential genes are analyzed by Enrichr program (https://maayanlab.cloud/Enrichr/) for pathway enrichment, followed by generation of bubble plot by using SRplot program (https://www.bioinformatics.com.cn/).

Then we determined the pathway enrichment in each cluster as shown (**Fig. 3d**). *Pdx1* was highly expressed in clusters 1, 2, 4 and 6, while *Pax6* was expressed in all clusters except cluster 5 (**Fig. 3n, o**). However, *Sst* was expressed in all clusters albeit the expression levels were lower in clusters 3 and 5 (**Fig. 3p**). The top 100 genes upregulated in cluster 1 were enriched with mTOR pathway, cluster 2 with ribosome biogenesis, RNA transport and one carbon pool by folate, cluster 3 with Ras signaling, cluster 4 and 5 with protein processing in endoplasmic reticulum, ribosome, oxidative phosphorylation and phagosome, cluster 6 with DNA repair and cell cycle (**Fig. 3q**). These data reflect the tumor cell heterogeneity.

### Conditional c-MYC induction by a Cre with improved coding sequence in the Dbh locus (Dbh-iCre) leads to development of neuroblastoma

A previous study utilized a *Dbh-iCre* mouse strain to cross *LSL-MYCN* mouse and successfully developed a genetic neuroblastoma model^26^. The improved coding sequence of Cre (iCre), followed by the bovine growth hormone polyadenylation signal, was introduced in-frame with the ATG of the *Dbh* gene^76^. We therefore surmised that *Dbh-iCre* might have a better recombination capacity to remove the floxed STOP cassette of the *c-MYC* gene. Because the *Dbh-iCre* strain has a mixed genetic background, we backcrossed the *Dbh-iCre* mice with the C57BL/6 stain for 10 generations before utilizing it to breed with the *CAG-MYC* mouse. We generated neuroblastoma models in genetically pure C57BL/6 background mice for several reasons. First, the mixed genetic background may affect tumorigenesis due to genetic drift occurring within the colony over time. Second, generating syngeneic mouse models using tumors from mixed genetic background mice is challenging because it will cause graft rejection and/or graft vs host disease upon transplantation. Syngeneic mouse models are valuable tools that can be used to quickly test therapies, particularly immunotherapies. Third, most of our knowledge of murine immunology and immunotherapies came from the studies of C57BL/6 mice. Thus, it may enhance the data reproducibility when interpreting the phenotypes and mechanisms related to the immune impact on tumorigenesis and therapy in C57BL/6 mouse models. Interestingly, the *CAG-MYC* mice bred with the *Dbh-Cre* (C57BL/6 background) progeny did not develop tumors; however, a high rate (16 out of 17 in one year) of tumorigenesis occurred when the *CAG-MYC* mice were bred with the *Dbh-iCre* mice in a C57BL/6 background (**Fig. 4a-c**). The tumors in *Dbh-iCre;CAG-MYC* mice were located in the abdomen (n=12), thoracic cavity (n=2) and neck (n=2), consistent with the previous genetic models showing that neuroblastoma occurred along the sympathetic chain such as superior cervical ganglion, adrenals or celiac ganglion^22,26^. In contrast, the *Dbh-iCre;LSL-MYCN* mouse model in a C57BL/6 background had a lower disease penetrance (5 out of 9), and tumors were located in the abdomen (**Fig. 4b, 4c**).

**Figure 4.**
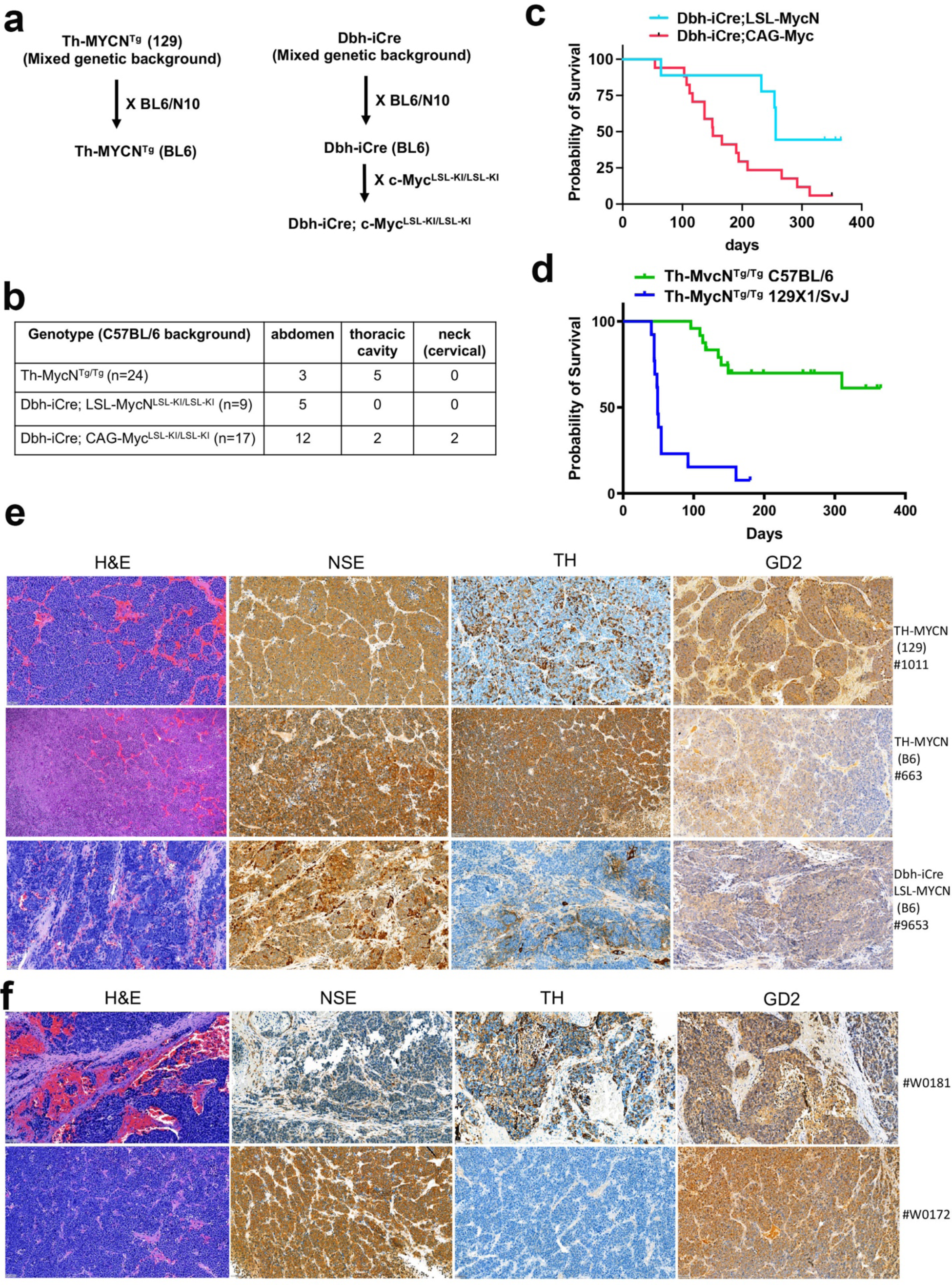
Neuroblastomas developed in Dbh-iCre;CAG-MYC mice. **a.** A schematic view of the generation of Th-MYCN^Tg^ and Dbh-iCre;CAG-MYC transgenic mice in C57BL/6 background. **b.** Spontaneous tumor development among Th-MYCN^Tg/Tg^, Dbh-iCre;LSL-MYCN^LSL-KI/LSL-KI^ and Dbh-iCre;CAG-MYC^LSL-KI/LSL-KI^ transgenic mice in a range of anatomic locations. **c.** Mouse survival curves of Dbh-iCre;LSL-MYCN^LSL-KI/LSL-KI^ and Dbh-iCre;CAG-MYC^LSL-KI/LSL-KI^ transgenic mice in 12 months. The death of mouse indicates the diagnosis of spontaneous tumor formation. **d.** Survival curves of mice with the homologous Th-MYCN transgene in C57BL/6 and 129X1/SvJ backgrounds in 12 months. **e.** Immunohistochemistry staining of tumor sections with the indicated markers for Th-MYCN^Tg/Tg^, Dbh-iCre;LSL-MYCN^LSL-KI/LSL-KI^ models. Scale bar = 50μm. **f.** Immunohistochemistry staining of tumor sections with the indicated markers for Dbh-iCre;CAG-MYC^LSL-KI/LSL-KI^ models. Scale bar = 50μm.

The *Th-MYCN* transgenic model was the first neuroblastoma GEMM, and has a much higher disease penetrance in a 129X1/ScJ background than the FVB/N or C57BL/6 background^22,77^. While the exact reason remains unknown, one early study revealed multiple secondary tumor susceptibility loci that may account for the tumor penetrance difference between the 129X1/ScJ background and other mouse backgrounds^77^. Here we also included this model by backcrossing it to the C57BL/6 strain for 10 generations (**Fig. 4a**). As expected, the 129X1/ScJ mice with homologous *Th-MYCN* transgene had an earlier disease onset and higher penetrance than in the C57BL/6 background. Eight out of 24 *Th-MYCN* C57BL/6 mice developed tumors in abdomen and thoracic cavity (**Fig. 4b, 4d**).

H&E staining showed that all tumors generated by *Th-MYCN* in 129X1/ScJ or C57BL/6, and *Dbh-iCre;LSL-MYCN* and *Dbh-iCre;CAG-MYC* in C57BL/6 mice showed primitive small, round blue cells typical for murine neuroblastomas (**Fig. 4e, f**), and were immunopositive for NSE. However, some tumors driven under the *Dbh-iCre* were TH-negative (**Fig. 4e, f**). Considering that *Th-Cre;CAG-MYC* mice developed PNETs, these data indicate that the expression of TH and DBH may not always overlap temporally and spatially during development. Nevertheless, all tumors showed strong positivity for disialoganglioside GD2 (**Fig. 4e, f**), a membrane surface maker in neuroblastoma and a therapeutic target for anti-GD2 immunotherapy or CAR T/NKT for treating neuroblastoma.

### Dbh-iCre;CAG-MYC tumors but not the Th-Cre;CAG-MYC tumors resemble human neuroblastoma

To determine the similarities between *Dbh-iCre;CAG-MYC* tumors and human neuroblastoma and *MYCN*-driven murine neuroblastomas, we compared the transcriptomics of tumors from our mouse models by RNA-seq with those of 15 different types of human pediatric solid tumors. We included a human neuroblastoma gene signature and the signature from a mouse model driven by *Th-MYCN/ALK^F1178L^* that were reported recently^24^. After identifying the gene signature for each type of the pediatric solid tumors sequenced at St Jude (one disease vs other diseases), we then used these 16 tumor-specific signatures (15 human cancers, 1 murine neuroblastoma) as gene sets to calculate the enrichment score for each mouse sample/human disease for clustering and heatmaps (**Fig. 5a-c**). Based on the ssGSEA results, we obtained 3 major clusters of all combined tumors (**Fig. 5a**). The cluster 2 was composed of human neuroblastoma, retinoblastoma, and all mouse tumors we generated, suggesting that these neuronally related tumors had a congruity than other types of tumors. However, the PNET driven by *Th-Cre;CAG-MYC* was segregated from other MYC-driven neuroblastomas including the one driven by *Dbh-iCre;CAG-MYC*, which branched into different subclusters (**Fig. 5a**). Then we further looked into the details of these tumors by comparing the human (**Fig. 5b**) and murine (**Fig. 5c**) neuroblastoma signatures with their transcriptomics. While to some degree, the PNET tumors shared some genes with human and murine neuroblastomas, which was not surprising, PNET tumors did not express *Phox2a* and *Phox2b*, two key neuroblastoma signature genes. However, the *Dbh-iCre;CAG-MYC* tumors and other mouse neuroblastoma models shared nearly identical gene signatures and were more closely related to the human neuroblastoma (**Fig. 5b, 5c**). Taken together, our data indicate that the *Dbh-iCre;CAG-MYC* tumors were neuroblastoma.

**Figure 5.**
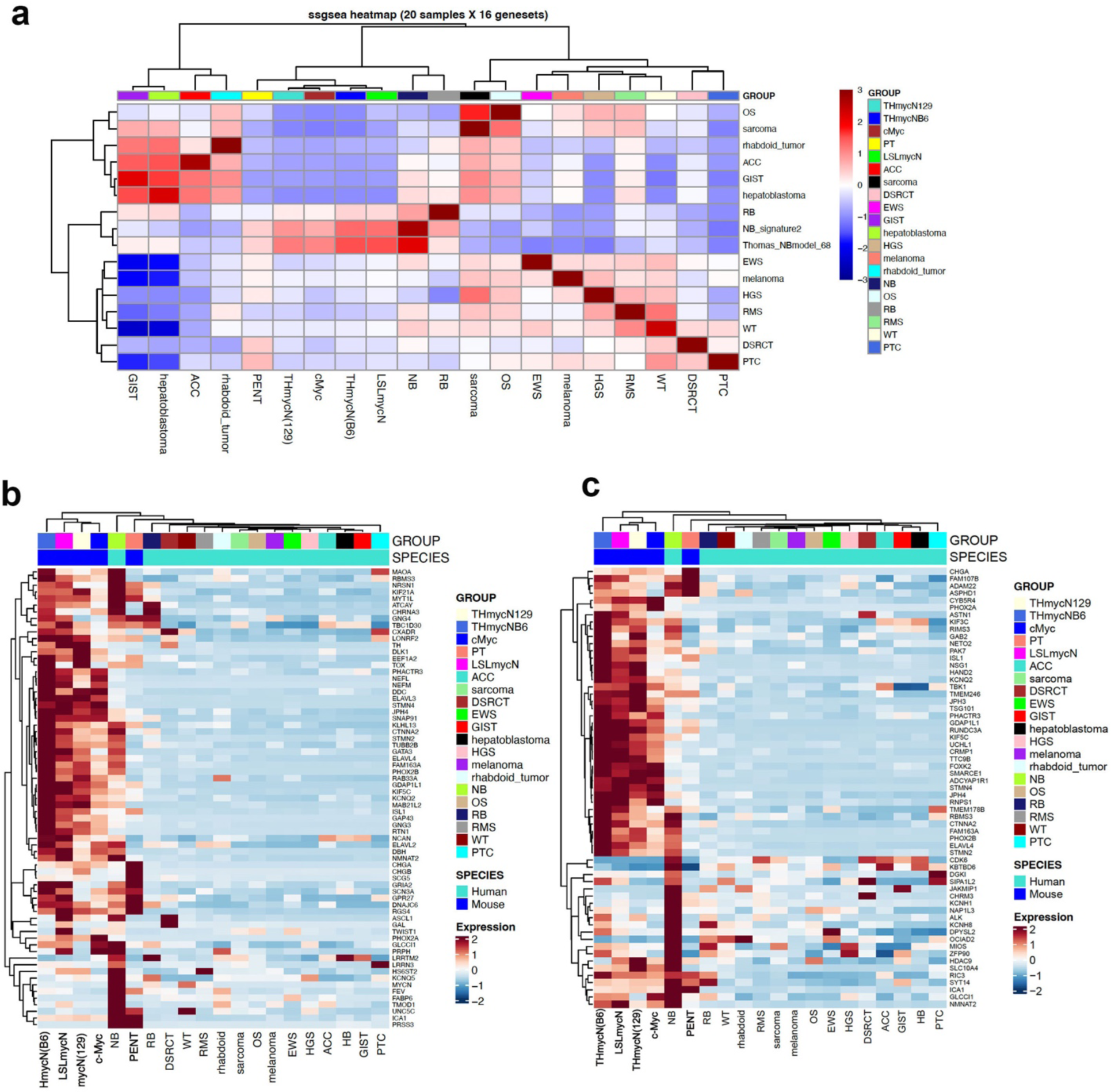
Transcriptomic similarity of Dbh-iCre;CAG-MYC tumors with neuroblastoma. **a.** ssGSEA heatmap showing dendrogram clusters of genetic mouse models and 14 human pediatric solid tumor types. OS = osteosarcoma, ACC = adrenocortical carcinoma, GIST =gastrointestinal stromal tumor, RB = retinoblastoma. NB = neuroblastoma, EWS = Ewing sarcoma, HGS =high grade sarcoma, RMS = rhabdomyosarcoma, WT = Wilm’s tumor, DSRCT = desmoplastic small round cell tumor, PTC = papillary thyroid cancer, PNET = pancreatic neuroendocrine tumor **b.** Heatmap based normalized gene expression showing that genetic neuroblastoma models generated in this study including Dbh-iCre;CAG-MYC tumors are clustered with human neuroblastoma. Pseudo-color indicates z-score of FPKM (Fragments Per Kilobase of transcript per Million mapped reads). **c.** Heatmap based normalized gene expression showing that genetic neuroblastoma models generated in this study including Dbh-iCre;CAG-MYC tumors are clustered with mouse Th-MYCN/ALK^F1178L^ neuroblastoma. Pseudo-color indicates z-score of FPKM.

### Mouse-derived allograft neuroblastoma models were established in immune-competent C57BL/6 host mice to test immunotherapies and targeted therapies

One advantage of GEMM in a C57BL/6 background is that it could allow us to generate syngeneic models by implanting tumor cells orthotopically and subcutaneously. To test whether the tumor cells from our GEMMs could form tumors through allograft implantation, we extracted single tumor cells in suspension and injected 5-8 million cells per mouse subcutaneously (**Fig. 6a**). The results showed that all tested tumor cells from the *Dbh-iCre;CAG-MYC*, *Th-MYCN*, and *Dbh-iCre;LSL-MYCN* models developed allograft tumors in C57BL/6 mice (**Fig. 6b-d**). While the tumor growth rate was variable in each mouse for each genetic tumor model, tumor volume reached over 2000mm^3^ within two months for the *Dbh-iCre;CAG-MYC* and *Dbh-iCre;LSL-MYCN* allograft models, and 4 months for the *Th-MYCN* allograft model. These data demonstrate that all these syngeneic models can be utilized to test therapies as they provide a reasonable therapeutic window based on the tumor growth rates.

**Figure 6.**
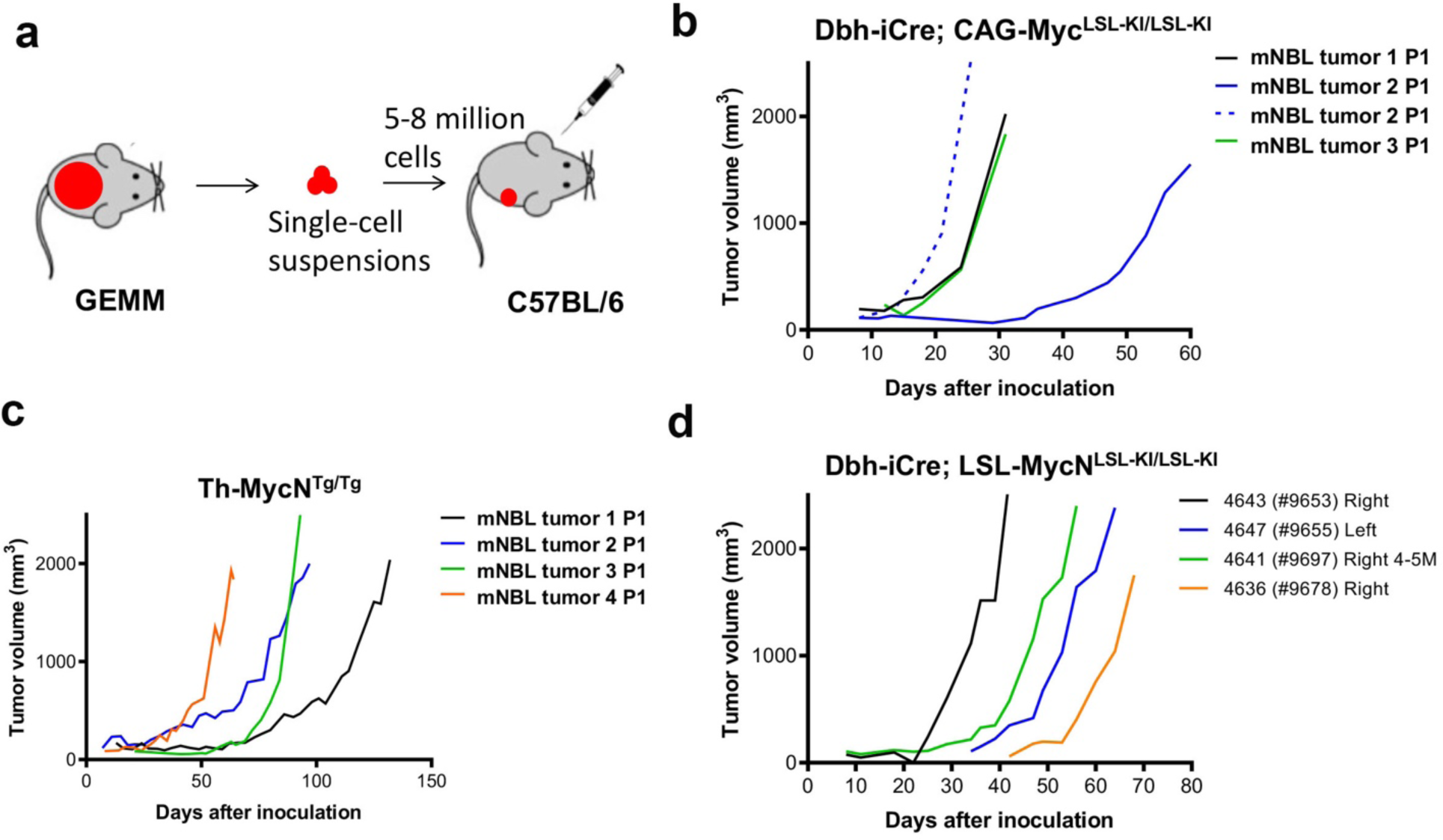
Establishing mouse derived allograft neuroblastoma models in immune competent C57BL/6 host mice. **a.** An overview of the establishment of mouse derived allograft neuroblastoma models by implanting 5-8 million single tumor cells in suspension from GEMM to immune competent C57BL/6 host mice. **b-d**. Tumor growth curves after the engraftment of tumor cells from GEMM Dbh-iCre;CAG-MYC^LSL-KI/LSL-KI^, Th-MYCN^Tg/Tg^, and Dbh-iCre;LSL-MYCN^LSL-KI/LSL-KI^ in C57BL/6 mice.

The application of anti-GD2 immunotherapy has greatly improved the survival of neuroblastoma patients when it is combined with a differentiating agent and chemotherapy^78–80^. Anti-GD2 immunotherapy has become a part of standard of care for high-risk neuroblastomas. Recent advances in developing GD2-based CAR T and CAR NKT therapies have provided additional evidence showing the potential of anti-GD2 immunotherapy for patients with relapsed disease^41,44^. The *Th-MYCN* mouse model in the 129X1/SvJ background was responsive to anti-GD2 immunotherapy^81^. Here we focused on the *Dbh-iCre;CAG-MYC* syngeneic models and tested their responses to immunotherapies and targeted therapies. First, we tested if this model could respond to the anti-GD2 immunotherapy. Before subcutaneous implantation, we confirmed that the *Dbh-iCre;CAG-MYC* tumor cells were GD2-positive by flow cytometry analysis, in which the human neuroblastoma cell line Lan-1 served as a positive control (**Fig. 7a**). One day after tumor cell inoculation, mice were treated with 200μg of anti-GD2 monoclonal antibody, twice weekly (**Fig. 7b**), and tumor growth was monitored (**Fig. 7c**). In comparison with the isotype IgG control group, the tumor growth was significantly delayed by the anti-GD2 therapy (**Fig. 7c**), which significantly extended mouse survival (**Fig. 7d**). These data demonstrated that our *c-MYC* syngeneic neuroblastoma model is suitable for testing the anti-GD2 immunotherapy. Then, we tested if this model could respond to the immune checkpoint blockade. It is known that neuroblastoma is immune “cold” due to a low mutation rate. Following a similar therapeutic schedule as used for anti-GD2 immunotherapy, the combination of anti-PD1 and anti-PD-L1 antibodies modestly but significantly delayed *c-MYC*-driven tumor growth (**Fig. 7e, 7f**). Recently, the U.S. Food and Drug Administration (FDA) approved eflornithine (Iwilfin, DFMO), an irreversible inhibitor of ornithine decarboxylase (ODC1), to reduce the risk of relapse in adult and pediatric patients with high-risk neuroblastoma who have demonstrated at least a partial response to prior multiagent, multimodality therapy, including anti-GD2 immunotherapy^46^. The rationale for targeting ODC1 activity for treating neuroblastoma is that the high-risk neuroblastoma is basically a MYC-driven cancer, and *ODC1* is a direct transcriptional target of MYC^47,48^ and a critical determinant of MYCN-mediated oncogenesis^47^. To test if our c-MYC neuroblastoma syngeneic model could respond to the DFMO treatment, we gave the mice with drinking water containing different concentrations of DFMO (**Fig. 7g**). While the low concentrations of DFMO (2g/L and 5g/L) showed no effect on tumor growth, 10g/L of DFMO in drinking water significantly delayed tumor growth (**Fig. 7h**). A combination of low concentrations of DFMO (2g/L) with either anti-GD2 or anti-PD1/PD-L1 did not significantly improve the anti-tumor effect (**Supplementary** Figure 8), probably due to the pharmacokinetics of DFMO at a low concentration. Taken together, our data demonstrated that the *Dbh-iCre;CAG-MYC* allograft model is suitable for testing immunotherapy and targeted therapy in neuroblastoma studies.

**Figure 7.**
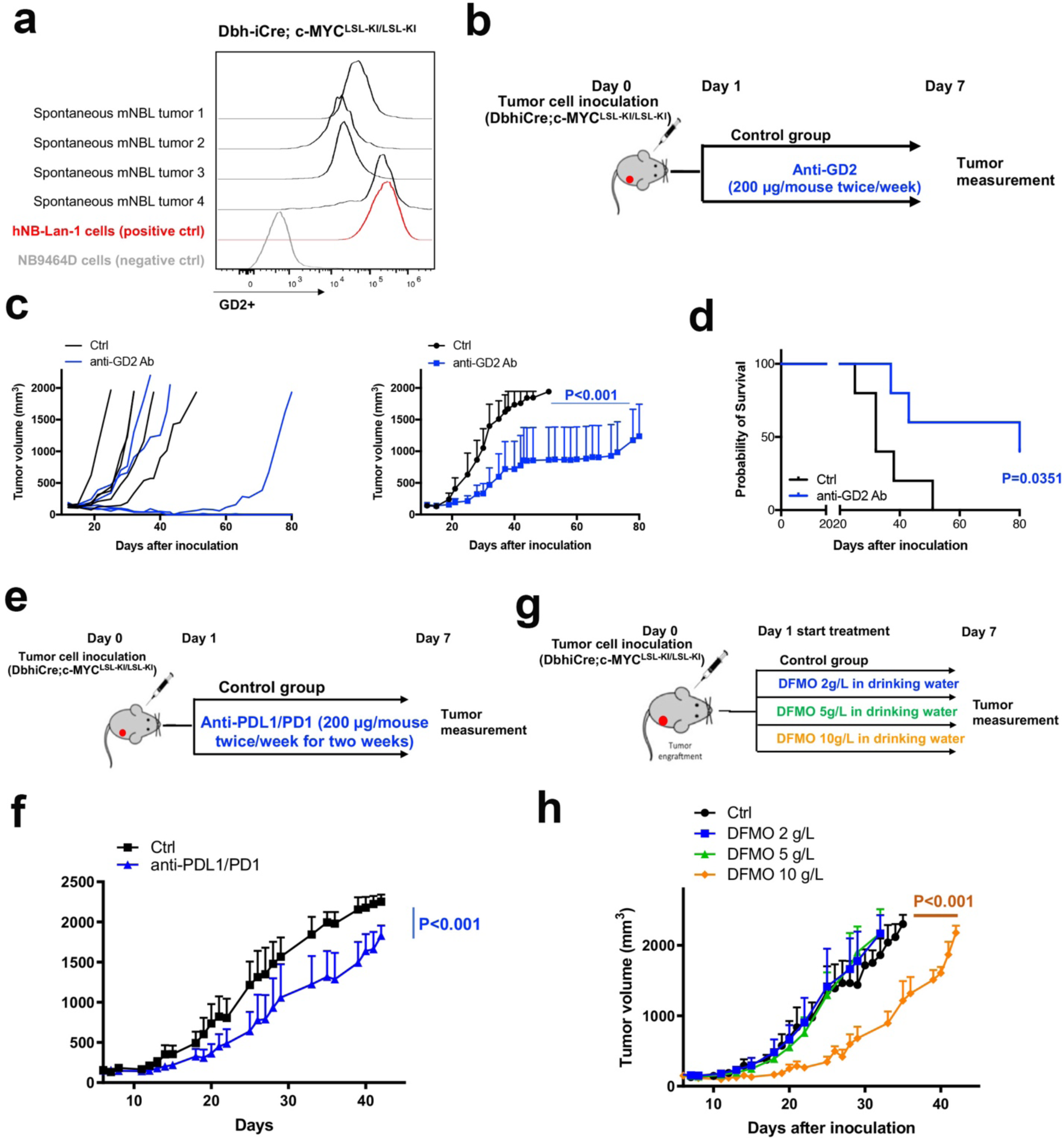
Immunotherapies and targeted therapies tested in mouse derived allograft neuroblastoma models. **a.** GD2 expression in Dbh-iCre;CAG-MYC tumor cells were confirmed by flow cytometry analysis. **b.** A schematic view of the experiment to test the response of mouse derived allograft models established with tumor cells from Dbh-iCre;CAG-MYC^LSL-KI/LSL-KI^ transgenic mice to anti-GD2 immunotherapy. **c.** Individual tumor growth curves (left panel) and average tumor growth curves (right panel) after syngeneic Dbh-iCre;CAG-MYC^LSL-KI/LSL-KI^ mice treated with IgG control (200 μg, intraperitoneal (i.p.) injection twice per week), or anti-GD2 antibody (200 μg, i.p. twice per week). Data represent mean ± s.d. (*n* = 5). *P* < 0.001 for IgG control versus anti-GD2 Ab by Wilcoxon test. **d.** Kaplan-Meier survival analysis of the c-MYC neuroblastoma models treated with IgG control or anti-GD2 Ab treatment. P=0.0062 for IgG control versus anti-GD2 Ab by Mantel-Cox test. **e.** A schematic view of c-MYC mouse derived allograft model for treatment with immune checkpoint blockade anti-PD1/PDL1 monoclonal antibody. **f.** Tumor growth curves after the engraftment of Dbh-iCre;CAG-MYC^LSL-KI/LSL-KI^ transgenic mice tumor cells to B57CL/6 mice, followed by the treatment of IgG control (200 μg, intraperitoneal (i.p.) injection twice per week), or anti-PDL1 antibody (200 μg, i.p. twice per week) and anti-PD1 antibody (200 μg, i.p. twice per week). Data represent mean ± s.d. (*n* = 5). *P* < 0.001 for IgG control versus anti-PDL1/PD1 Ab by Wilcoxon test. **g.** A schematic view of the experiment to test the response of c-MYC neuroblastoma model to DFMO treatment. **h.** c-MYC neuroblastoma models were treated with regular drinking water, or different doses of DFMO (2g/L, 5g/L, 10g/L) in drinking water. The tumor volumes of control group and DFMO-treated groups were monitored daily. Data represent mean ± s.d. (*n* = 3). *P* < 0.001 for control versus 10g/L DFMO by Wilcoxon test.

## Discussion

In contrast to cancer cell line xenograft models, GEMMs develop *de novo* tumors in a natural immune-proficient microenvironment, capture both tumor cell-intrinsic and cell-extrinsic factors that drive tumor initiation and progression, and display tumor heterogeneity^82^. Designing and interpreting the neoplasms that arise in GEMMs is another strategy for modeling the spectrum of diseases occurring in human patients. These models may also be reliable surrogates for patients that help discover novel therapies for disease intervention or prevention, dissect the impact of the tumor microenvironment, and evaluate mechanisms of drug resistance. It has been over 25 years since the development of the first neuroblastoma GEMM generated by the *MYCN* oncogene^22^. While *MYCN* amplification occurs in 50% of high-risk neuroblastoma, the c-MYC activity is also deregulated in neuroblastoma, whose expression is mutually exclusive to the *MYCN* amplification. Thus, a c-MYC–based GEMM has been sought in the neuroblastoma field to validate the oncogenic drivers of neuroblastoma and enable the testing of novel therapeutic approaches to provide unexplored translational opportunities. In this study, we successfully developed a c-MYC-driven neuroblastoma GEMM after having tried several different mouse lines that express Cre recombinase in neural crest progenies, the cell of origin of neuroblastoma. Tumors arising in *Dbh-iCre;CAG-MYC* closely mimic the histopathological and molecular features of human neuroblastoma, and respond to the FDA-approved immunotherapy and targeted therapy for neuroblastoma treatment. So far, all successful genetic neuroblastoma models have been based on the activation of MYC in either *Th*-or *Dbh*-positive cells^22–28^. Unexpectedly, we found that crossing of *Dbh-Cre* with *CAG-MYC* was unable to initiate tumorigenesis, suggesting that the Cre with improved coding sequences (iCre) has a better activity to drive c-MYC-mediated tumorigenesis. This finding might be important to mouse genetics and developmental biology as the iCre may server as a better Cre driver. In a C57BL/6 background, mice with *Dbh-iCre;CAG-MYC* showed a better disease penetrance than those with *Dbh-iCre;LSL-MYCN*. While the reason for this difference was not clear, it could be due to the differences of biological functions of MYCN and c-MYC or MYCN and c-MYC strains.

Interestingly, c-MYC activation in *Th-Cre* expressing cells induced tumorigenesis in pancreatic islets with a low penetrance in older mice, and the tumor spectrum was different from neuroblastoma. The differences in disease onset time, anatomical locations, histopathology and transcriptomics between the *Th-Cre;CAG-MYC* and *Dbh-iCre;CAG-MYC* mice indicate that temporal and spatial activation of c-MYC determines the disease entities and spectrum of neoplasia that may arise in these models. One early study revealed that, during differentiation of sympathetic neurons in chick embryos, the *Th* and *Dbh* mRNAs appeared at the same developmental period, but later *Dbh* can be detected in both *Th*-positive and –negative cells^83^. Genetic reporters to label the *Th*– and *Dbh*-positive cells in mice revealed that, while both *Th* and *Dbh* were expressed in sympathetic neurons, it was *Th* but not *Dbh* that was expressed in pancreatic islets^84,85^. One independent study showed that Th expression occurs in both β cells and δ cells of mouse islets^86^. In the mouse pancreas at embryonic day E12.5–E13.5, ∼10% of early glucagon^+^ cells expressed *Th*, and the number of insulin^+^ cells in the *Th ^−/−^* embryonic pancreas was decreased^87^, suggesting that TH may play an important role in regulating differentiation of islet cells. Knowledge from these early development biology studies provided a potential explanation why *Th-Cre;CAG-MYC* mice developed PNETs with molecular features of δ cells that expressed high levels of *Sst* that encodes somatostatin. Somatostatinoma is an extremely rare neuroendocrine tumor with an incidence of 1 in 40 million individuals and accounts for less than 5% of PNETs. The tumor originates from the δ cells of the pancreas and predominantly contains somatostatin with trace quantities of other pancreatic hormones such as insulin, glucagon, gastrin, and vasoactive intestinal polypeptide^88^. This tumor type may be observed in conjunction with hereditary cancer predisposition syndromes including familial endocrine and neural tumor syndromes like neurofibromatosis 1 (NF1) and von Hippel Lindau syndromes^89,90^. Therefore, the *Th-Cre*;CAG-MYC model may provide a unique tool for studying the role of MYC in inherited rare cancers.

A recent transcriptomic and proteomic study of human PNET revealed 4 subtypes: proliferative, PDX1-high, α cell-like and stromal/mesenchymal. The PDX1-high group barely had genetic mutations of PNET genes such as *MEN1* and *ATRX*^91^. The *Th-Cre;CAG-MYC* somatostatinoma tumors express high levels of *Pdx1*, suggesting that they might resemble the human PDX1-high PNET. Since *ATRX* was found to be mutated in both human neuroblastomas and PNET, we attempted to knock out *Atrx* in our model system to see if loss of ATRX function would promote tumorigenesis, which was not the case (data not shown).

The expression of the *MYCN* transgene by the *Th* promoter led to neuroblastoma development. Why c-MYC activation in a *Th-Cre* line developed a rare PNET but not neuroblastoma was puzzling. There could be many reasons that caused distinct tumor spectra in these models including the ways which oncogenes were activated and differences between the expression of MYCN and c-MYC in development, which is known to affect ontogeny, promote stemness and self-renewal, and altering or blocking differentiation pathways^92^. Nevertheless, we have created two unique genetic models: c-MYC-driven neuroblastoma and somatostatinoma. The c-MYC and MYCN neuroblastoma models that we generated are transplantable to C57BL/6 mice with 100% success. These models can serve as useful tools and sources in the neuroblastoma field to study cancer initiation, progression and tumor heterogeneity, test new targeted and immunotherapies, and understand mechanisms of tumor microenvironment and therapy resistance.

## Methods

### Animals

All experiments that involved the use of mice were performed in accordance with the guidelines outlined by the Nationwide Children’s Hospital (Dr. Ruoning Wang) and St Jude Children’s Research Hospital (Dr. Jun Yang) Institutional Animal Care and Use Committee (IACUC). Mice were housed with ambient temperature and humidity with a 12 h light /12 h dark cycle controlled under specific-pathogen-free conditions (SPF) at the Nationwide Children’s Hospital and St Jude Children’s Research Hospital mouse facility. Mice were allowed to feed and drink ad libitum. The maximal tumor burden permitted was 20% of mouse body weight, and in our experiments, the maximal tumor burden was not exceeded. Transgenic mice were euthanized through CO_2_ inhalation with 3 liters/min in the mouse cage and followed by cervical dislocation when moribund or determined by a veterinarian in the Animal Research Center at Nationwide Children’s Hospital and St Jude. For therapy studies in subcutaneous xenograft mouse models, the mice were euthanized through CO_2_ inhalation with 3 liters/min in the mouse cage and followed by cervical dislocation when the tumor volume reached 2000 mm^3^ or mice became moribund.

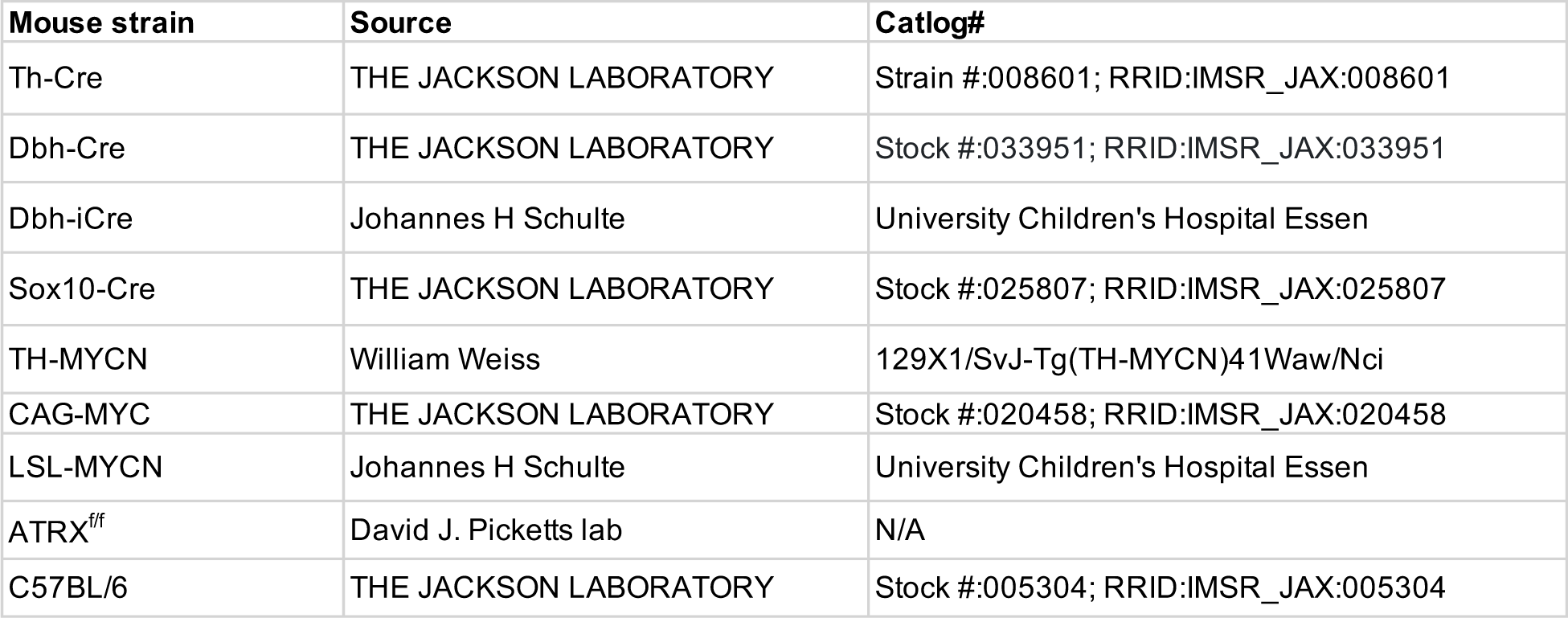

## Mouse strains and sources

### Mouse genotyping

DNA was extracted from ear clips using Thermo Scientific Phire Tissue Direct PCR Master Mix kit according to the manufacturer’s protocol (Thermo Fisher Scientific). The PCR was performed in a total reaction volume of 10 µL containing 0.5 µM of each primer, 100 ng of DNA and 2X Phire Tissue Direct PCR Master Mix or DreamTaq Green 2X master mix (Thermo Fisher Scientific). PCR products were separated by 2-3% agarose gel electrophoresis, stained with ethidium bromide and photographed under UV-light. TH-MYCN transgene genotype was determined by PCR genotyping according to the previously described method^93^ with the following primers: Chr18F1: 5′-ACTAATTCTCCTCTCTCTGCCAGTATTTGC-3′, Chr18R2: 5′-TGCCTTATCCAAAATATAAATGCCCAGCAG-3′, and OUT1: 5′-TTGGCACACACAAATGTATATACACAATGG-3′. Primers used for detecting TH-Cre were oIMR1084 (5′-GCGGTCTGGCAGTAAAAACTATC-3′), oIMR1085 (5′-GTGAAACAGCATTGCTGTCACTT-3′), oIMR7338 (5′-CTAGGCCACAGAATTGAAAGA TCT-3′), and oIMR7339 (5′-GTAGGTGGAAATTCTAGCATCATCC-3′) according to the genotyping method described by the Jackson Laboratory. Primers used for detecting Dbh-iCre were Dbh-iCre-Fwd: 5′-CTGCCAGGGACATGGCCAGG –3′ and Dbh-iCre-Rev: 5′-GCACAGTCGAGGCTGATCAGC –3′ according to the previously described method^26^. For genotyping LSL-MYCN mice, PCR was performed with the following primers, LSL-MYCN-Fwd: 5’-ACCACAAGGCCCTCAGTACC-3’ and LSL-MYCN-Rev: 5’-TGGGACGCACAGTGATGG –3’ to confirm LSL-MYCN transgene integration, and primers 5’-CTCTTCCCTCGTGATCTGCAACTCC –3’ and 5’-CATGTCTTTAATCTACCTCGATGG –3’ to validate the presence of a wild-type allele at the ROSA26 locus as previously described^26^. CAG-MYC transgene genotype was determined by PCR genotyping using primers 13840: 5’-CCAAAGTCGCTCTGAGTTGTTATC –3’ and 13841: 5’-GAGCGGGAGAAATGGATATG –3’ for wild-type allele, and 16298: 5’-CCAAGAGGGTCAAGTTGGA –3’, and oIMR8191: 5’-GCAATATGGTGGAAAATAAC –3’ for CAG-MYC transgene integration according to the Jackson Laboratory protocol. Primers for detecting Dbh-Cre were based on the Jackson Laboratory listed (Dbh1: 5′-AGG CAT AAA TGG CAG AGT GG-3′, Dbh2: 5′-CAT GTC CAT CAG GTT CTT GC-3′, Dbh3:5′-TGG AGC TGG AAG TGG ATG AT-3′). Sox10-Cre genotype was determined by PCR genotyping using primers (SOX10Cre-IPCF: 5’-GAC AAA ATG GTG AAG GTC GG-3’, SOX10Cre-IPCR: 5’-CAA AGG CGG AGT TAC CAG AG-3’, Sox10Cre-F: 5’-CAC CTA GGG TCT GGC ATG TG-3’, Sox10Cre-R: 5’-AGG CAA ATT TTG GTG TAC GG-3’). For ATRX flox/flox mice, these primers were used to determine heterogeneity (1kb and 1.5kb two bands) and homozygosity (only ∼1.5kb band) (ATRX-F: 5’-GGTTTTAGATGAAAATGAAGAG –3’, ATRX-R1: 5’-TGAACCTGGGGACTTCTTTG –3’, ATRX-R2: 5’-CCACCATGATATTCGGCAAG –3’).

### Cell lines

Lan-1 cell line was kindly provided by Dr. Xiaotong Song. NB975A2, a mouse neuroblastoma cell line derived from spontaneous tumors of TH-MYCN mice, was a gift from Dr. Rimas Orentas. All NBL cells used were grown in Dulbecco’s modified Eagle’s medium with 10% fetal calf serum in a 37 °C humidified atmosphere of 95% air and 5% CO_2_. Lan-1 was validated by short tandem repeat (STR) profiling using PowerPlex® 16 HS System (Promega). A polymerase chain reaction (PCR)-based method was used to screen for mycoplasma employing the LookOut® Mycoplasma PCR Detection Kit (MP0035, Sigma-Aldrich) and JumpStart™ Taq DNA Polymerase (D9307, Sigma-Aldrich) to ensure cells were free of mycoplasma contamination.

### Magnetic resonance imaging

Magnetic resonance imaging (MRI) was performed on a Bruker Biospec 94/20 MRI system (Bruker Biospin MRI GmbH, Ettlingen, Germany). Prior to scanning, mice were anesthetized in a chamber (3% Isoflurane in oxygen delivered at 0.5 L/min) and maintained using nose-cone delivery (1-2% Isoflurane in oxygen delivered at 0.2 L/min). Animals were provided thermal support using a heated bed with warm water circulation and a physiological monitoring system to monitor breath rate. MRI was acquired with a mouse brain surface receive coil positioned over the mouse head and placed inside an 86 mm transmit/receive coil. After the localizer, T2-weighted Rapid Acquisition with Refocused Echoes (RARE) sequences were performed in the coronal (TR/TE = 2000/20.4 ms, matrix size = 256 x 256, field of view = 20 mm x 20 mm, slice thickness = 0.5 mm, number of slices = 16) and axial (TR/TE = 2500/23 ms, matrix size = 256 x 256, field of view = 20 mm x 20 mm, slice thickness = 0.5 mm, number of slices = 32) orientations.

### Ultrasound imaging of mouse models

Fur was removed from the ventral side of each animal using Nair. Technicians in the St. Jude Center for In Vivo Imaging and Therapeutics performed ultrasound scanning on mice weekly using VEVO-3100 (FUJIFILM VisualSonics, Toronto, Canada) and determined tumor volumes using VevoLAB 5.7.1 software. All ultrasound data were acquired in a blinded fashion.

### GD2 flow cytometry analysis

The tumors were harvested as a single-cell suspension, washed with PBS, and blocked with blocking buffer. Subsequently, cells were incubated with APC-conjugated anti-GD2 (Biolegend, Catalog # 357305) or APC-conjugated anti-mouse IgG (Biolegend, Catalog # 402205) (1 μg per 10^6^ cell) master mix for 1 hour on ice. Following this, cells were washed twice with PBS supplemented with 5% FBS. Cells were then spun down and resuspended in PBS for analysis. Flow cytometry data were acquired on Novocyte (ACEA Biosciences) and signal overlay analysis was conducted using FlowJo software.

### Establishment of mouse-derived allografts for therapies

Isolation and preparation of single cell suspension from tumor samples were conducted as follows. Spontaneous tumor tissues with a volume of around 2000 mm^3^ were dissected from Dbh-iCre;CAG-MYC transgenic mice, washed with phosphate-buffered saline (PBS) in a sterile 35mm dish. Tumor samples were then minced in 5mL DMEM (Corning) supplemented with 10% fetal calf serum and 1% penicillin–streptomycin, and transferred to a gentleMACS C tube (gentleMACS™ Octo Dissociator) and digested with collagenase IV(100U/mL) and DNase I (10μg/mL) on the gentleMACS™ Octo Dissociator (gentleMACS™ Octo Dissociator) under program “37C_h_TDK_1”. The resulting cell mixture was filtered through a sterile 70uM cell strainer, centrifuged, and washed with PBS. The cells were then treated with red blood cell (RBC) lysis buffer (Biolegend) to remove debris and centrifuged to collect primary neuroblastoma cells. After counting the tumor cells, the cell suspension solution was centrifuged, and the cell pellet was incubated on ice for 5 min. Cells were then resuspended with 30% volume of ice-cold PBS and 70% volume of MatriGel (Corning). To establish the mouse-derived allograft model, 8-week-old C57BL/6 mice (gender-matched with the transgenic mice bearing primary tumor) were inoculated with 3-8 × 10^6^ tumor cells subcutaneously into the flanks. In all the allograft experiments, immunotherapy with InVivoMAb Rat IgG2b Isotype control (BioXcell, Cat# BE0090) or indicated antibodies anti-GD2 (BioXCell BE0318), anti-PD1 (BioXCell, Catalog #BE0146) and anti-PDL1 (BioXCell, Catalog #BE0101) (200 μg/mouse) was performed i.p. twice per week, and DFMO (Eflornithine HCl monohydrate) (Carbosynth LLC, Cat# FE11666) treatment was administered (2 g/L) in drinking water bottle to the indicated experimental groups until the end of the experiment. For all the animal studies, tumor sizes were measured daily by caliper 7 days after implantation, and tumor volume was calculated by length × width^2^ × (π / 6). For survival studies, animals were monitored daily until the endpoint was reached, that is when the experimental animals had (1) tumor diameter that reached 2 cm; (2) tumor ulceration reaching 1 cm in diameter, or persistent bleeding as a result of ulceration; (3) a 20% loss of body weight; (4) breathing difficulty or (5) poor mobility.

### Pathology and Immunohistochemistry

Tumors confirmed by imaging were harvested and immersed in 10% neutral-buffered formalin for at least 72 hours. After fixation, tissues were embedded in paraffin, sectioned at 4-μm, mounted on positively charged glass slides (Superfrost Plus, Thermo Fisher Scientific), and dried at 60°C for 20 minutes before dewaxing and staining with hematoxylin and eosin (Richard-Allan Scientific) or used for immunohistochemistry. Coverslips were placed using the HistoCore SPECTRA Workstation (Lecia Biosystems). Pathology evaluation was conducted blindly to any experimental conditions. Hematoxylin & eosin (HE) and serial immunohistochemistry (IHC) sections were scanned to a 20x scalable magnification using a PANNORAMIC 250 PIII slide scanner (3DHistech, Ltd.) and imaged using HALO v3.6.4134.137 (Indica Labs).

A Ventana Discovery Ultra autostainer (Roche, Indianapolis, IN) and the following conditions were used to detect Tyrosine hydroxylase (TH, Millipore, AB152, 1:500), Vimentin (ABCAM, ab92547, 1:500), S100 (DAKO, Z031129, 1:2000), Glucagon (Cell Signaling, 2760, 1:100), Insulin (Lifespan Biosciences, LS-B2526, 1:800), Cytokeratin Oscar (Covance, SIG-3465-1000, 1:250), Chromogranin A (ImmunoStar, 20085, 1:1500), and Synaptophysin (ABCAM, ab32127, 1:400): Heat-induced epitope retrieval, Cell Conditioning Solution ULTRA CC1 (950-224, Roche) and visualization with DISCOVERY OmniMap anti-rabbit HRP (760-4311), DISCOVERY ChromoMap DAB kit (760-159). A Ventana DISCOVERY ULTRA automated stainer (Ventana Medical Systems, Inc., Tucson, AZ) and the following conditions were used to detect neuron-specific enolase (NSE, DAKO, M087301, 1:300): Heat-induced epitope retrieval (HIER), Cell conditioning media 2 (CC2), 32 minutes at 37°C; Visualization with rabbit anti-mouse secondary at 1:500 for 16 minutes (Abcam, ab133469), DISCOVERY OmniMap anti-rabbit HRP (760-4311), and DISCOVERY ChromoMap DAB kit (760-159). An IntelliPath FLX autostainer (Biocare medical) and the following conditions were used to detect Cytokeratin AE1/AE3 (Millipore, MAB3412, 1:10,000): Proteinase K treatment for 10 minutes (DAKO, Carpinteria CA, S3020), UltraVision Quanto Mouse on Mouse HRP kit (Thermo Fischer, Freemont, CA, TL-060-QHDM), UltraVision Quanto Detection System HRP DAB kit (Thermo Fischer, Freemont, CA, TL-060-QHD) and Mayer’s hematoxylin stain (Thermo Fischer, Fremont, CA, TA-125-MH). An IntelliPath FLX autostainer and the following conditions were used to detect 14.G2a anti-GD2 disialoganglioside antibody (1:500 dilution; BD Biosciences, catalog no. 554272, 1:50): HIER with Citrate pH 6.0 at 110°C (Invitrogen, 005000), UltraVision Quanto Mouse on Mouse HRP kit (Thermo Fischer, Freemont, CA, TL-060-QHDM), UltraVision Quanto Detection System HRP DAB kit (Thermo Fischer, Freemont, CA, TL-060-QHD) and Mayer’s hematoxylin stain (Thermo Fischer, Fremont, CA, TA-125-MH).

### RNA extraction and RNA-seq and analysis

In order to obtain high-quality and intact RNA from the normal pancreas, the pancreas was perfused with the RNase inhibitor solution RNAlater (Qiagen, 76106) before isolation from the littermate control mice. The isolated pancreas with perfusion was cut into small pieces for total RNA extraction. Total RNAs from normal pancreas and tumor tissues were performed using the RNeasy Mini Kit (Qiagen, Cat#74136) according to the manufacturer’s instructions.

Total stranded RNA sequencing data were processed by the internal AutoMapper pipeline. Briefly the raw reads were first trimmed (Trim Galore version 0.60), mapped to mouse genome assembly (GRCm38, mm10) (STAR v2.7) and then the gene level values were quantified (RSEM v1.31) based on GENCODE annotation (VM22). Low count genes were removed from analysis using a CPM cutoff corresponding to a count of 10 reads in at least one sample group and only confidently annotated (level 1 and 2 gene annotation) and protein-coding genes are used for differential expression analysis. Normalization factors were generated using the TMM method, counts were then transformed using voom and transformed counts were analyzed using the lmFit and eBayes functions (R limma package version 3.42.2). The significantly up– and down-regulated genes were defined by at least 2-fold changes and adjusted p-value < 0.05. Then gene set enrichment analysis (GSEA) was carried out using gene-level log2 fold changes from differential expression results against gene sets in the Molecular Signatures Database (MSigDB 6.2) (gsea2 version 2.2.3). GSEA parameters (number of permutations =1000, permutation type = gene_set, metric for ranking genes = Signal2Noise, Enrichment statistic = Weighted).

### Single nuclear RNA-seq data and processing

Single nuclear RNA-seq data was generated from a frozen tumor harvested from mouse neuroendocrine model. Single nuclei were isolated and subjected to RNA-sequencing following the protocol of Slyper et al.2020. Briefly, we adopted TST method (Tween with Salts and Tris) ^94^. The tissue was chopped on ice for 10 min on TST buffer, filtered through 40µm cell strainer before pelleting the cells. The cell pellet was then washed twice in the ST buffer followed by filtering nucleus again in a 20μm strainer. The nuclei were counted using the Luna cell counter, 8000 nuclei of the single cell suspension were loaded onto the 10X chromium Chips – V3.1-Single index of Single cell gene expression kit. The cDNA and library were sequenced on Novaseq following 10X Genomics recommended depth and raw data were processed through Cell Ranger Software (10X Genomics). Clusters with distinct transcriptomic signatures were derived using the LCA method ^95^. Differentially expressed genes were extracted using the NBID algorithm^96^ with the following filters: FDR ≤ 0.05, fold change ≥ 2^0.5^, and average TPM in the higher cluster ≥ 500. The graphs were presented by using Cloupe (10xGenomics).

### Single sample GSEA

The RNA-seq data of 14 pediatric solid tumor types (CSTN_RNAseq_data_FPKM) were obtained from St Jude (https://www.stjude.org/research/why-st-jude/data-tools/childhood-solid-tumor-network.html). Then RNA-seq data from our genetic neuroblastoma models and the 14 pediatric solid tumor types were combined, only keeping the common gene symbols between human and mouse. The signature for each disease (one disease vs other diseases) was identified by differential gene expression analysis to generate genesets to calculate the enrichment score for each mouse sample/human disease. We also included the signature of Th-MYC/ALK^F1178L^ neuroblastoma model^24^. Lastly, ssGSEA score was used for clustering and visualizing results in dendrogram and heatmap.

## Data availability

GEO accession number: GSE254964 (for sncRNA-seq) and GSE255205 (for bulk RNA-seq) To review GEO accession GSE255205: https://www.ncbi.nlm.nih.gov/geo/query/acc.cgi?&acc=GSE255205&token=mjkjomcapxyfxgz To review GEO accession GSE254964: https://www.ncbi.nlm.nih.gov/geo/query/acc.cgi?&acc=GSE254964&token=sjclkcsspvyxtcj

## Acknowledgements

We thank the staff in St. Jude Hartwell Center and Comparative Pathology Core for their dedication and expertise. We thank Dr. Johannes H Schulte from the University Children’s Hospital Essen for providing the Dbh-iCre and LSL-MYCN strains. This work was partly supported by V Foundation (V2014-001, R.W.), American Cancer Society-Research Scholar (130421-RSG-17-071-01-TBG, J.Y.;128436-RSG-15-180-01-LIB, R.W.), National Cancer Institute (1R01CA229739-01 and 1R01CA266600-01A1, J.Y.; U01CA232488, 2R01AI114581-06, R01CA247941 and 1R01AI175004-01, R.W.), Comprehensive Cancer Center core grant CA021765, and the American Lebanese Syrian Associated Charities (ALSAC). The content is solely the authors’ responsibility and does not necessarily represent the official views of the National Institutes of Health.

## Author contributions

T.W., L.L., J.F., S.N, and S.S. performed experiments and data collection. G.T., T.C., M.J., and J.S. performed small animal imaging. H.S. and M.L. performed pathological anatomy studies and immunohistochemistry analysis with help from L.H., N.Y., R.J., H.J. and X.C performed bioinformatics analysis. D.J.P provided resources. A.M., A.M.D., E.S.G., J.E., R.W., and J.Y were involved in project supervision and conceptualization. R.W. and J.Y. designed the experiments. J.Y. wrote the manuscript with help from all others.

