## Supplemental figures for "Conditional c-MYC activation in catecholaminergic cells drives distinct neuroendocrine tumors: neuroblastoma vs somatostatinoma"

### Supplementary Figure

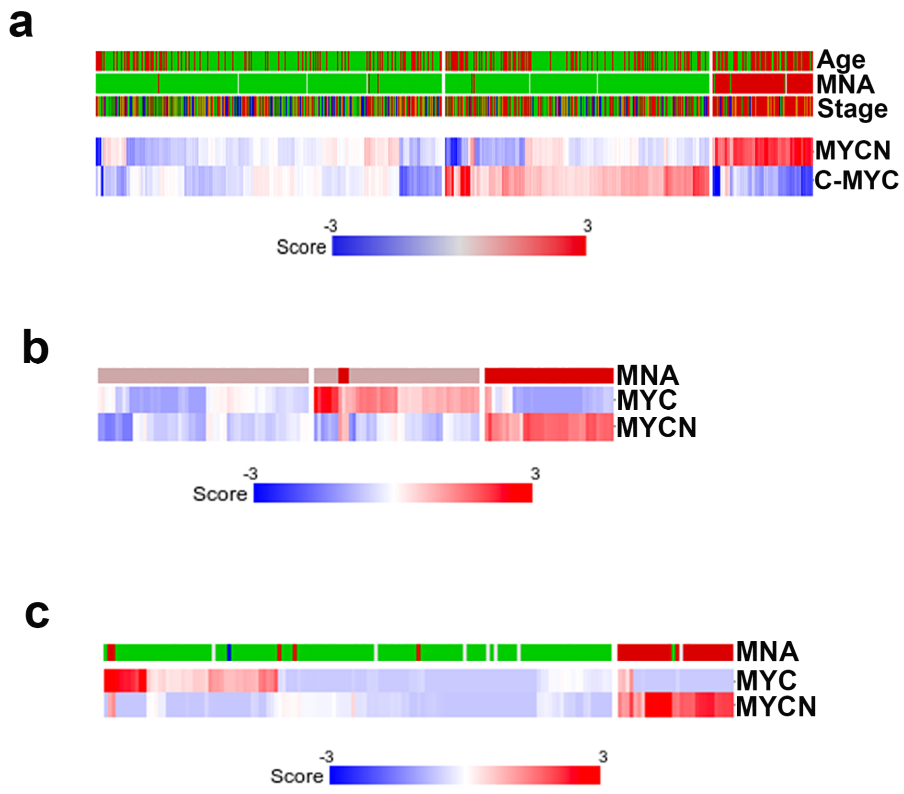


**Supplementary Figure 1. The expression of *MYCN* and *MYC* is mutually exclusive in human neuroblastomas. a**. Heatmap for expression of MYCN and MYC in neuroblastomas from Kocak dataset (GSE45547). Age (red > 18 months; green < 18 months); MNA (MYCN amplification, red = amplification; green = non-amplification); stage (red = stage 4; blue = stage 4S; brown = stage 3; dark green = stage 2, green = stage 1) **b**. Heatmap for expression of MYCN and MYC in neuroblastomas from Westermann dataset (GSE73517). Red=MNA, light purple = non-MNA. **c**. Heatmap for expression of MYCN and MYC in neuroblastomas from TARGET dataset (https://gdc.cancer.gov/about-data/publications#/?programs=TARGET&order=desc). MNA = MYCN amplification. Red =MNA, green =non-MNA. All heatmaps were made through online R2 program (https://hgserver1.amc.nl/cgi-bin/r2/main.cgi).

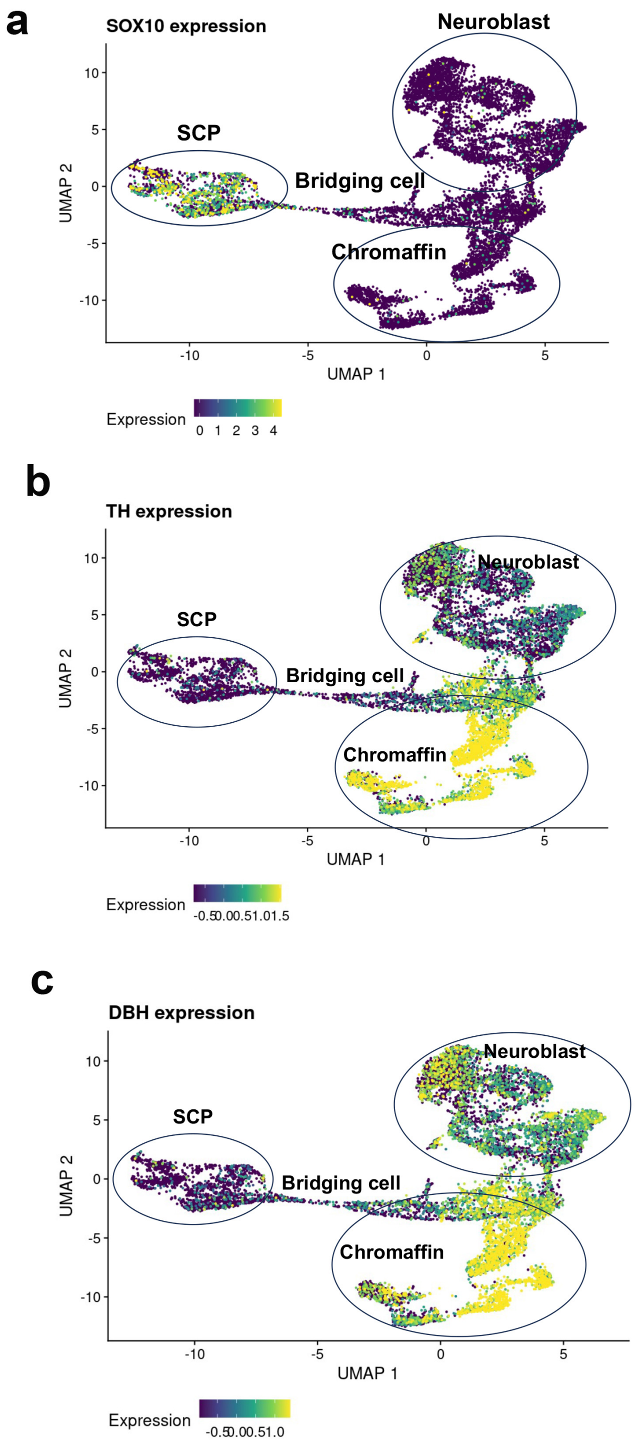


**Supplementary Figure 2. Single cell RNA expression of SOX10, TH and DBH in human adrenal medulla.** 4 clusters of cells including Schwann cell precursor (SCP), bridging cells, neuroblasts and chromaffin cells in human adrenal medulla, which are presumed to be the cell origin of neuroblastoma. Data were obtained from Jansky et al (https://adrenal.kitz-heidelberg.de/developmental_programs_NB_viz/).

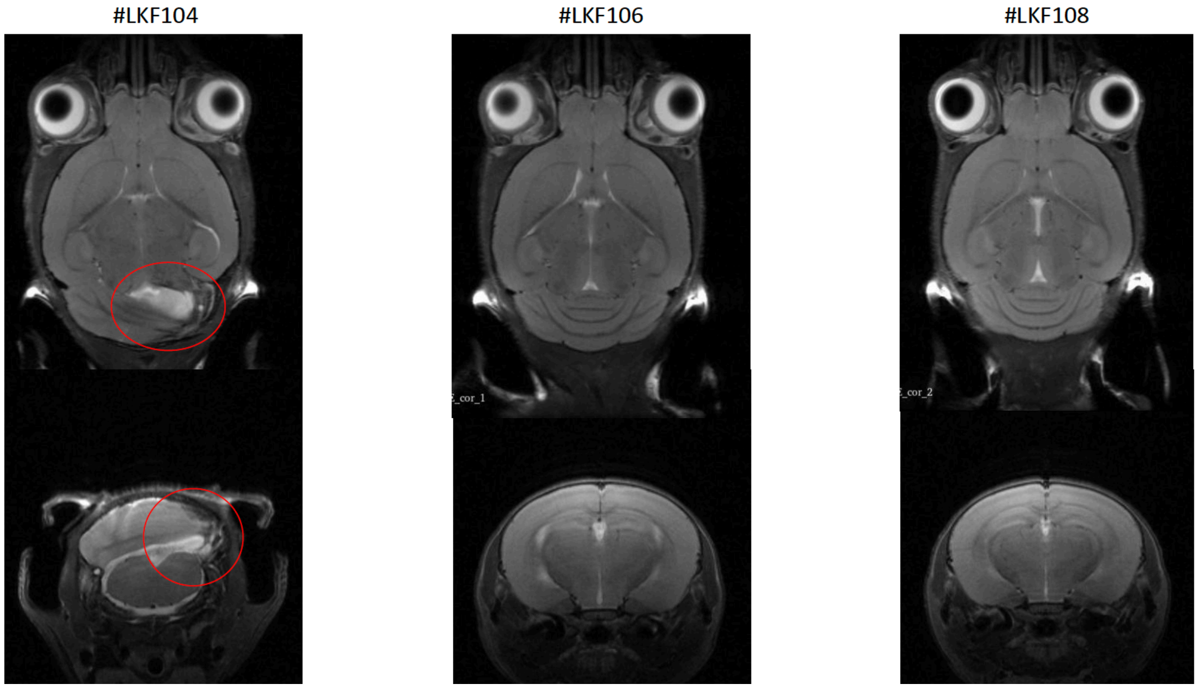


**Supplementary Figure 3. C-MYC activation by Sox10-Cre leads to brain lesions in young mice (Glioma).** Magnetic resonance imaging scan showed 1 out 3 mice had brain lesion.

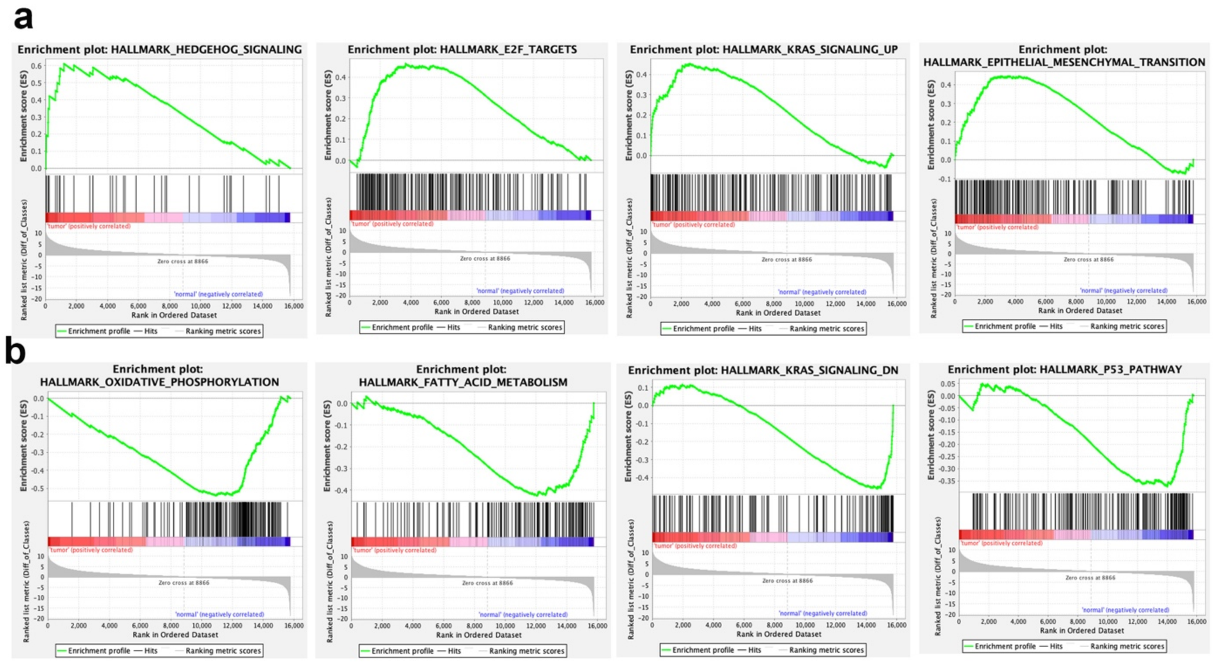


**Supplementary Figure 4. GSEA analysis of Th-Cre;CAG-MYC tumors vs age matching normal pancreatic tissues.** a. Genesets significantly downregulated in tumors (nominal P<0.01). b. Genesets significantly upregulated in tumors (nominal P<0.01).

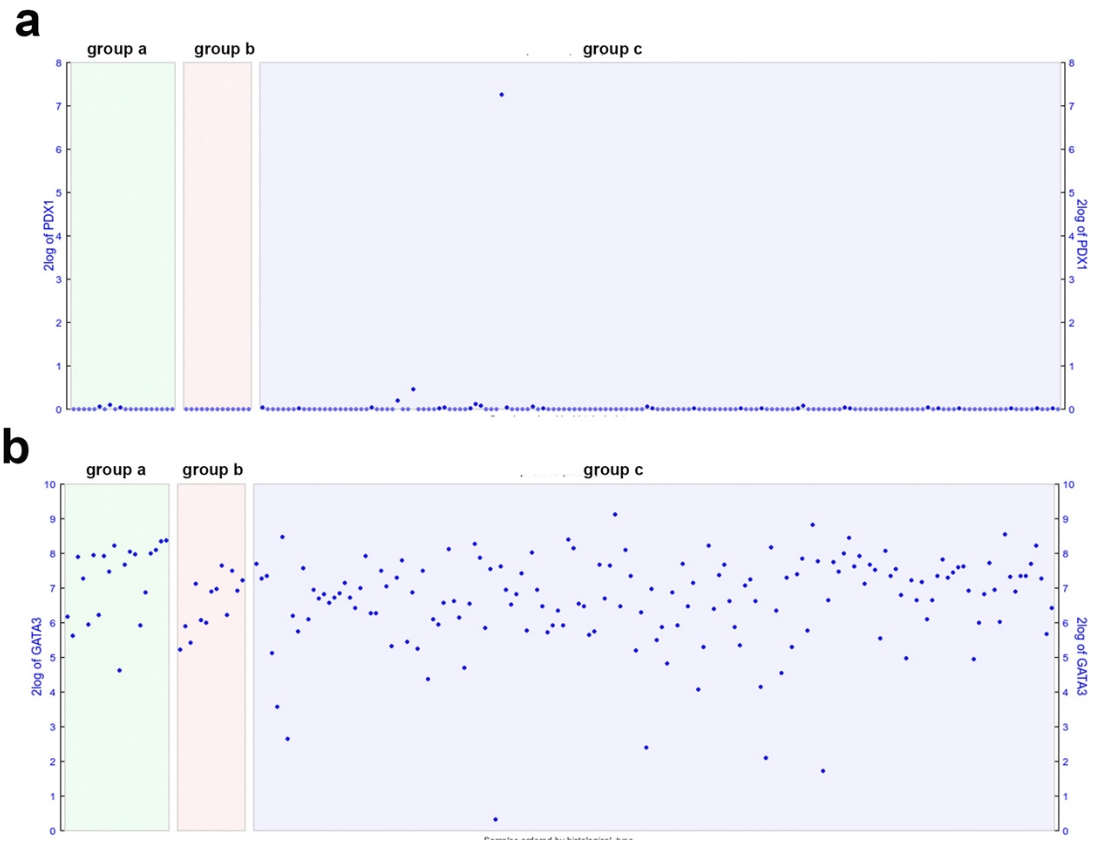


**Supplementary Figure 5. Expression of PDX1 and GATA3 in human paraganglioma and pheochromocytoma.** Group a = paraganglioma, group b = paraganglioma and extra-adrenal pheochromocytoma, group c = pheochromocytoma. Data were from TCGA (https://hgserver1.amc.nl/cgi-bin/r2/main.cgi).

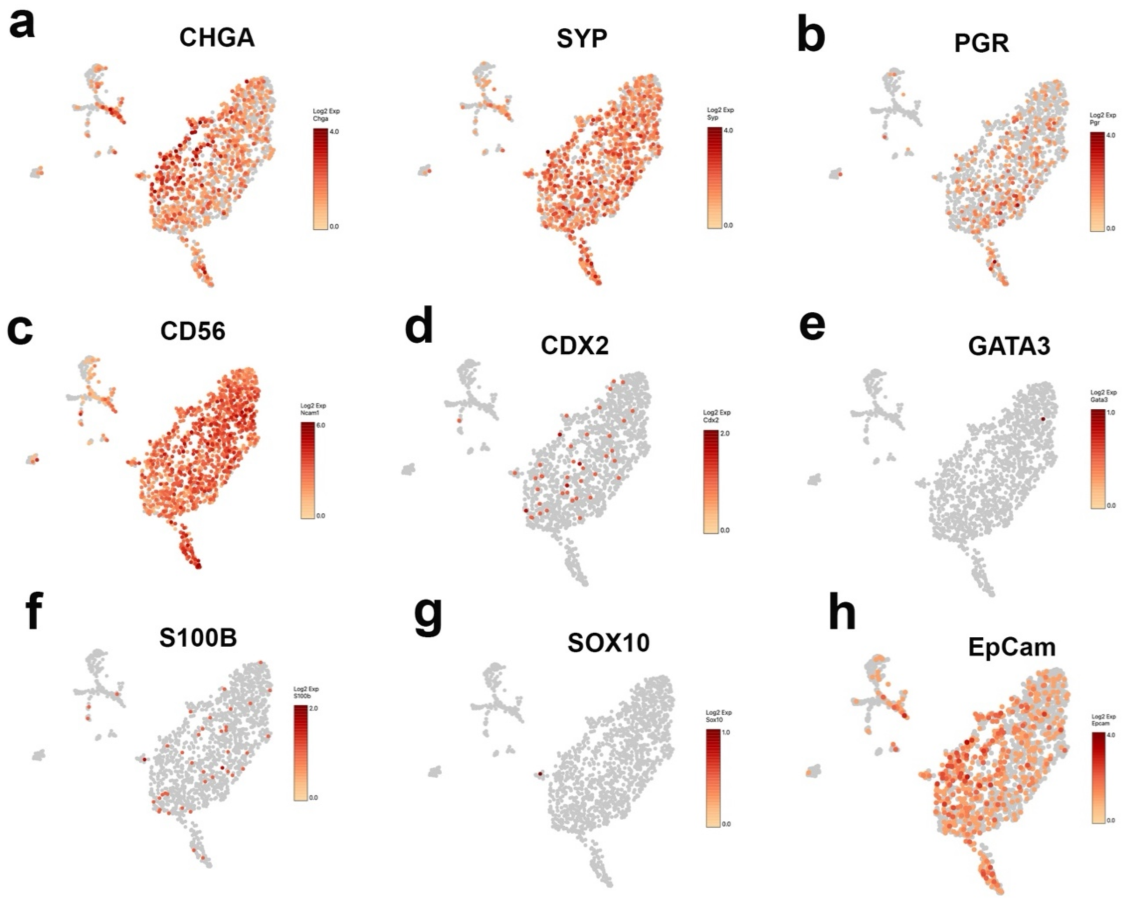


**Supplementary Figure 6. Diagnostic markers indicate the cell origin of Th-Cre;CAG-MYC tumors. a.** *Chga* and *Syp*, two markers for neuroendocrine tumors. **b**. *Pgr*, a marker for pancreatic neuroendocrine tumor. **c**. *CD56*, encoding neural cell adhesion molecule (NCAM), a marker for neuroendocrine tumors. **d**. *Cdx2,* a highly sensitive and specific marker of adenocarcinomas of intestinal origin. **e**, *Gata3*, a marker to differentiate pancreatic neuroendocrine tumors and paraganglioma and pheochromocytoma. ***f***. S100B, a marker could be expressed in pancreatic neuroendocrine tumors and paraganglioma and pheochromocytoma. **g**. Sox10, marker for Schwannian and melanocytic lineages and paraganglioma and pheochromocytoma. **h**. *EpCam*, a marker for epithelial cell origin of tumors.

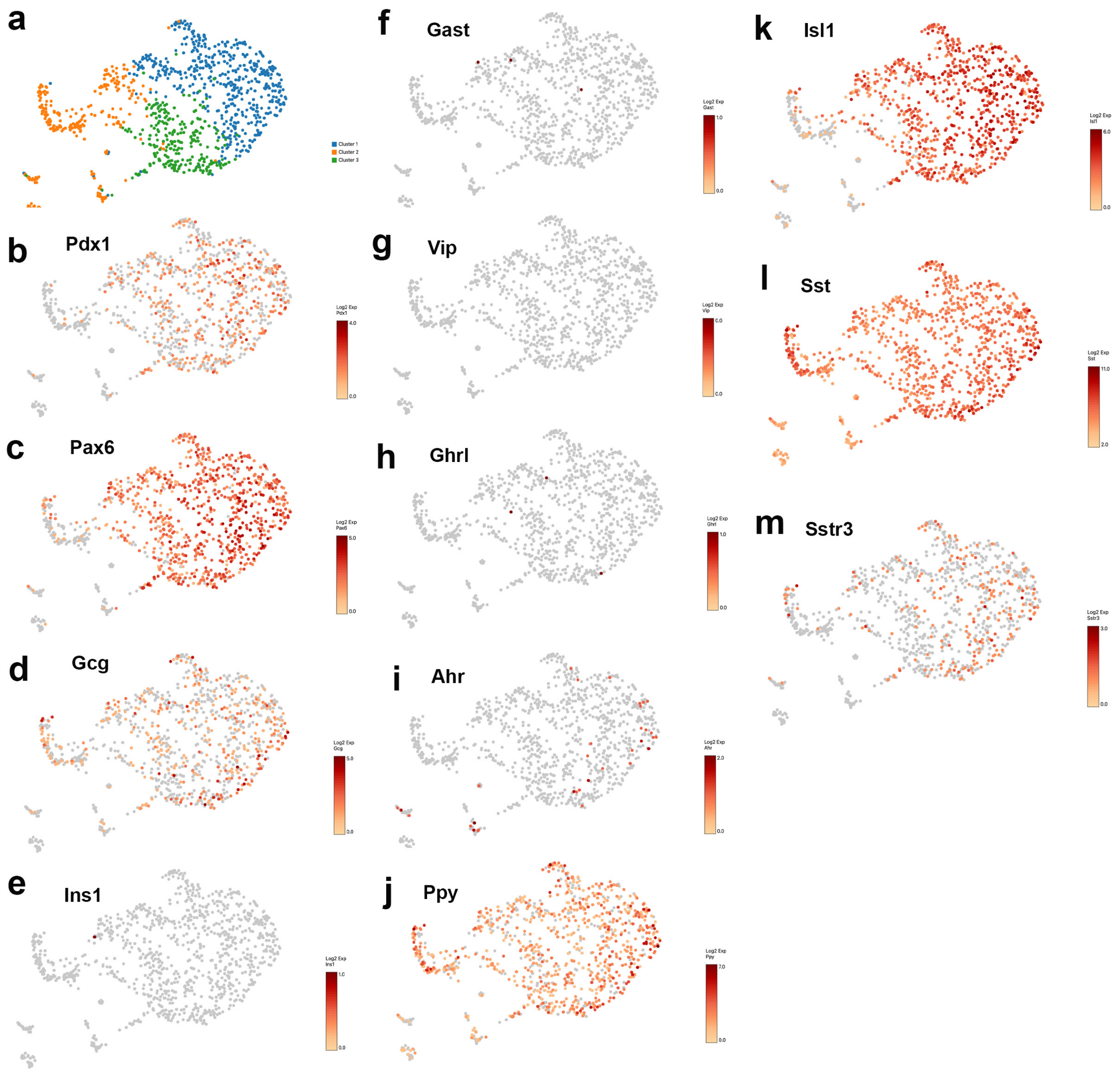


**Supplementary Figure 7. Th-Cre;CAG-MYC tumor cells express markers of somatostatinoma. a**. UMAP showing 3 clusters of tumor cells. **b, c**. Expression of Pdx1 and Pax6, the two transcriptional factors for development of pancreatic endocrine cells. d. *Gcg* (encoding glucagon, a marker for α cells and glucagonoma). **e**. *Ins1* (encoding insulin, a marker for β cells and insulinoma). **f**. *Gast* (encoding gastrin, a marker for G cells and gastrinoma). **g**. *Vip* (a marker for VIPoma). **h**. *Ghrl* (encoding ghrelin, a marker for e cells). **i, j.** *Ahr* and *Ppy* that encodes aryl hydrocarbon receptor and pancreatic polypeptide, markers for PP cells. **k-m**. *Isl1* (encoding ISL LIM homeobox 1, a δ cell transcriptional factor), *Sst* (encoding somatostatin, a marker for δ cells and somatostatinoma), *Sstr3* (encoding somatostatin receptor 3), markers for δ cell and somatostatinoma.

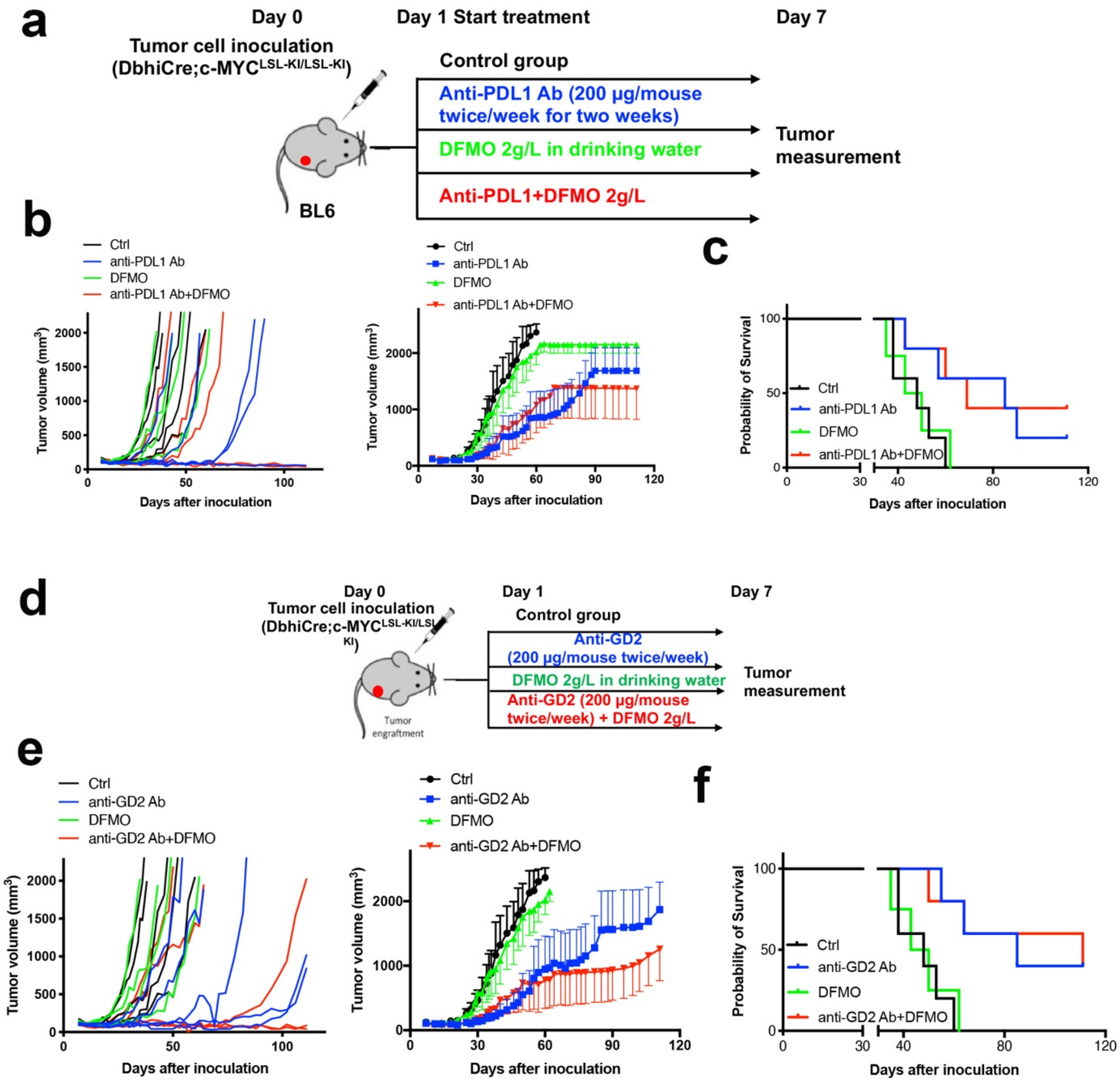


**Supplementary Figure 8. Combination of low dose of DFMO with anti-PDL1 or anti-GD2 is not superior. a.** A schematic view of the experiment to test the response of mouse derived allograft models established with tumor cells from Dbh-iCre;CAG-MYCLSL-KI/LSL-KI transgenic mice to combination of DFMO with anti-PDL1 immunotherapy. **b.** Individual tumor growth curves (left panel) and average tumor growth curves (right panel) after mice treated with IgG control or anti-PDL1 (200 µg, intraperitoneal (i.p.) injection twice per week), DFMO (2g/L in drinking water) or combination. Data represent mean ± s.d. ($n = 5$). $P < 0.001$ for IgG control versus anti-PDL1 by Wilcoxon test. **c**. Kaplan-Meier survival analysis of mice by Mantel-Cox test. **d**. A schematic view of c-MYC syngenetic mouse models for treatment with combination of DFMO with anti-GD2. Immunotherapy. **e.** Individual tumor growth curves (left panel) and average tumor growth curves (right panel) after mice treated with IgG control or anti-GD2 (200 µg, intraperitoneal (i.p.) injection twice per week), DFMO (2g/L in drinking water) or combination. Data represent mean ± s.d. ($n = 5$). $P < 0.001$ for IgG control versus anti-GD2 by Wilcoxon test. **c**. Kaplan-Meier survival analysis of mice by Mantel-Cox test. Please note, the control group and DFMO group in Figure S8b and 8e serve as the same control groups to compare with the combination therapies.

#### Table S1. The top 20 signaling pathways upregulated and downregulated in Th-MYC;CAG-MYC tumors.

| GENE SET (Tumor vs Normal) | NES | NOM p-val | FDR q-val |
|---|---|---|---|
| REACTOME_POTASSIUM CHANNELS | 2.15 | 0 | 0 |
| REACTOME_NEURONAL SYSTEM | 2.15 | 0 | 0 |
| KEGG_MEDICUS REFERENCE CA2 ENTRY VOLTAGE GATED CA2 CHANNEL | 2.04 | 0 | 0.001 |
| REACTOME_INWARDLY RECTIFYING K CHANNELS | 2.03 | 0.001 | 0.001 |
| REACTOME_PROTEIN PROTEIN INTERACTIONS AT SYNAPSES | 2.03 | 0 | 0.001 |
| REACTOME_RESOLUTION OF_SISTER CHROMATID COHESION | 1.98 | 0 | 0.002 |
| REACTOME_NEUROTRANSMITTER RECEPTORS AND POSTSYNAPTIC SIGNAL TRANSMISSION | 1.96 | 0 | 0.003 |
| REACTOME_KINESINS | 1.95 | 0 | 0.003 |
| REACTOME_TRANSMISSION ACROSS CHEMICAL SYNAPSES | 1.95 | 0 | 0.003 |
| REACTOME_NEUREXINS AND NEUROLIGINS | 1.95 | 0 | 0.003 |
| REACTOME_REGULATION OF INSULIN SECRETION | 1.9 | 0 | 0.008 |
| BIOCARTA_LIS1 PATHWAY | 1.9 | 0 | 0.007 |
| REACTOME_GABA B RECEPTOR ACTIVATION | 1.89 | 0.001 | 0.007 |
| KEGG_MEDICUS VARIANT MUTATION CAUSED ABERRANT HTT TO RETROGRADE AXONAL TRANSPORT | 1.89 | 0 | 0.006 |
| REACTOME_L1CAM INTERACTIONS | 1.89 | 0 | 0.006 |
| KEGG_MEDICUS REFERENCE CARDIAC TYPE VGCC RYR SIGNALING | 1.89 | 0 | 0.006 |
| REACTOME_TRAFFICKING OF AMPA RECEPTORS | 1.88 | 0.001 | 0.007 |
| REACTOME_UPTAKE AND ACTIONS OF BACTERIAL TOXINS | 1.88 | 0 | 0.008 |
| PID_AURORA B PATHWAY | 1.87 | 0 | 0.008 |
| KEGG_MEDICUS REFERENCE SKELETAL TYPE VGCC RYR SIGNALING | 1.86 | 0.001 | 0.008 |
| REACTOME_EUKARYOTIC TRANSLATION INITIATION | -2.88 | 0 | 0 |
| KEGG_MEDICUS REFERENCE TRANSLATION INITIATION | -2.84 | 0 | 0 |
| KEGG_RIBOSOME | -2.84 | 0 | 0 |
| REACTOME_SRP DEPENDENT COTRANSLATIONAL PROTEIN TARGETING TO MEMBRANE | -2.79 | 0 | 0 |
| REACTOME_RESPONSE OF EIF2AK4 GCN2 TO AMINO ACID DEFICIENCY | -2.71 | 0 | 0 |
| REACTOME SELENOAMINO ACID METABOLISM | -2.69 | 0 | 0 |
| WP_CYTOPLASMIC RIBOSOMAL PROTEINS | -2.68 | 0 | 0 |
| REACTOME_TRANSLATION | -2.63 | 0 | 0 |
| REACTOME_EUKARYOTIC TRANSLATION ELONGATION | -2.52 | 0 | 0 |
| REACTOME_NONSENSE MEDIATED DECAY NMD | -2.49 | 0 | 0 |
| REACTOME_ACTIVATION OF THE mRNA UPON BINDING OF THE CAP BINDING COMPLEX AND EIFS AND SUBSEQUENT BINDING TO 43S | -2.48 | 0 | 0 |
| REACTOME RESPIRATORY ELECTRON TRANSPORT | -2.47 | 0 | 0 |
| REACTOME_RESPIRATORY ELECTRON TRANSPORT ATP SYNTHESIS BY CHEMIOSMOTIC COUPLING AND HEAT PRODUCTION BY UNCOUPLING PROTEINS | -2.41 | 0 | 0 |
| KEGG_MEDICUS REFERENCE MITOCHONDRIAL COMPLEX UCP1 IN THERMOGENESIS | -2.4 | 0 | 0 |
| REACTOME_METABOLISM OF AMINO ACIDS AND DERIVATIVES | -2.39 | 0 | 0 |
| REACTOME_COMPLEX I BIOGENESIS | -2.38 | 0 | 0 |
| REACTOME_SARS COV 1 MODULATES HOST TRANSLATION MACHINERY | -2.38 | 0 | 0 |
| KEGG_MEDICUS PATHOGEN SARS COV 2 NSP1 TO TRANSLATION INITIATION | -2.35 | 0 | 0 |
| KEGG_MEDICUS REFERENCE ELECTRON TRANSFER IN COMPLEX I | -2.35 | 0 | 0 |
| KEGG_MEDICUS VARIANT MUTATION INACTIVATED PINK1 TO ELECTRON TRANSFER IN COMPLEX I | -2.35 | 0 | 0 |
